# Cooperative interactions among females and even more extraordinary sex ratios

**DOI:** 10.1101/2020.06.08.118786

**Authors:** Ryosuke Iritani, Stuart A West, Jun Abe

**Affiliations:** RIKEN Interdisciplinary Theoretical and Mathematical Sciences (iTHEMS), Wako, Saitama 351-0198, Japan; Department of Zoology, University of Oxford, Oxford, UK; Faculty of Liberal Arts, Meijigakuin University, 1518 Kamikurata-cho, Totsuka, Yokohama City, Kanagawa, 244-8539, Japan

**Keywords:** Kin selection, Local resource competition, Local mate competition, Local resource enhancement, Population structure

## Abstract

Hamilton’s local mate competition theory provided an explanation for extraordinary female biased sex ratios in a range of organisms. When mating takes place locally, in structured populations, a female biased sex ratio is favoured to reduce competition between related males, and to provide more mates for males. However, there are a number of wasp species where the sex ratios appear to more female biased than predicted by Hamilton’s theory. We investigated theoretically the extent to which cooperative interactions between related females can interact with local mate competition to favour even more female biased sex ratios. We found that: (i) cooperative interactions between females can lead to sex ratios that are more female biased than predicted by local competition theory alone; (ii) sex ratios can be more female biased when the cooperative interactions are offspring helping parents before dispersal, rather than cooperation between siblings after dispersal. Our results can be applied to a range of organisms, and provide an explanation for the extreme sex ratio biases that have been observed in *Sclerodermus* and *Melittobia* wasps.

## Introduction

Sex ratio theory has provided one of the most productive and successful areas of evolutionary biology (Charnov 1982; Hardy 2002; West 2009). Theory predicts a number of situations in which individuals are expected to adjust the sex of their offspring in response to local conditions (Charnov 1982; Frank 1998). This theory has been applied to explain variation in the offspring s ex ratio (proportion males) across a range of taxa, from malaria parasites to ants to birds (Bourke & Franks 1995; Hardy 2002; West 2009).

One of the major challenges is to explain when sex ratios are biased away equal investment in the sexes. Hamilton’s (1967) local mate competition (LMC) theory provides an explanation for female biased sex ratios observed in parasitic wasps (e.g., *Scelionidae*, Waage 1982; *Alfonsiella*, Greeff 2002; *Apanteles*, Tagawa 2000; Gu & Dorn 2003; and *Nasonia*, Werren 1983; Shuker *et al.* 2006; Burton-Chellew *et al.* 2008), aphids (e.g., *Prociphilus oriens*; Yamaguchi 1985), and a number of fig wasps (Herre 1985). Specifically, Hamilton showed that if *n* haplodiploid females lay eggs in a patch, and that mating occurs before females disperse, then the evolutionarily stable strategy (ESS; Maynard Smith & Price 1973) is to produce an offspring sex ratio of (*n*−1)/(2*n*) (Fig 1), which predicts female biased offspring sex ratios (smaller than 1/2) and becomes less biased as more females lay eggs in a patch (i.e., as *n* larger). Succeeding work (Taylor 1981; Bulmer 1986; Frank 1986; Taylor 1988, 1992; Frank 1998) made it clear that in Hamilton’s LMC theory, selection on the female bias is mediated by the balance among three factors: (i) a benefit for reduced competition between sons, (ii) a benefit for production of more mates (daughters) for those sons (“mating bonus”; Frank 1998), and (iii) a cost for stronger local resource competition among females (LRC; Clark 1978). Hamilton’s LMC theory has been extremely successful in explaining variation in the offspring sex ratio, both across species and between individuals (Taylor 1981, 1994; Avilés 1993; Gardner & West 2006; Shuker *et al.* 2004, 2006; Gardner *et al.* 2009; West 2009; Rodrigues & Gardner 2015).

**Figure 1:**
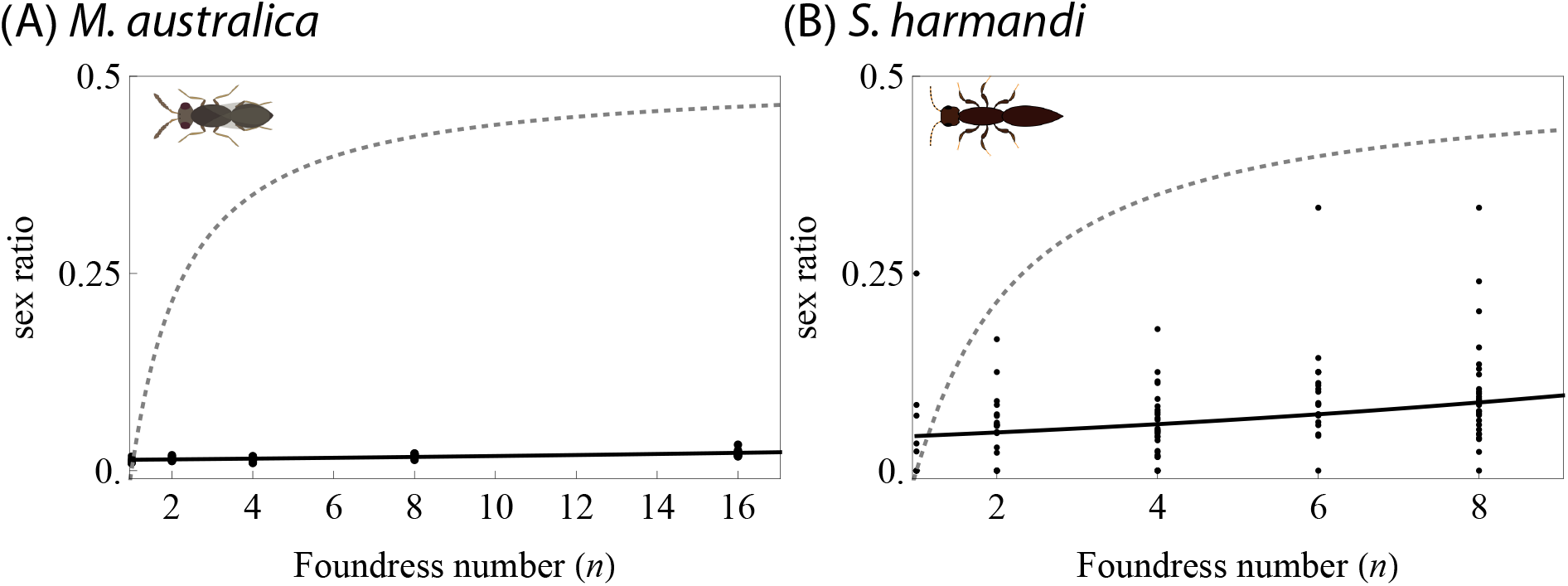
Extremely female-biased sex ratio in (A) *Melittobia australica* from Abe *et al.* (2003), and (B) *Sclerodermus harmandi* from Tang *et al.* (2014) and Kapranas *et al.* (2016). Both species haplodiploids. Outliers removed for (B), as in the original articles Tang *et al.* (2014) and Kapranas *et al.* (2016). Note that the horizontal axes are scaled differently. Dotted lines: Reference sex ratio give by ((*n* − 1)(4*n* − 2)/(2*n*(4*n* − 1)) (evolutionarily stable sex ratio for haplodiploids with *d*_f_ = 1). Solid line in (A): predicted values by generalized linear models; in (B): shown in Tang *et al.* (2014).

However, there are a number of cases where females produce extremely female biased offspring sex ratios, which do not appear to be completely explained by LMC theory. One example is provided by *Melittobia* wasps, where females of several species produce approximately 2 % male offspring when ovipositing alone (*n* = 1), and hardly change their offspring sex ratio when more females lay eggs on a patch (larger; Fig 1A; Abe *et al.* 2003, 2014). Another example is provided by *Sclerodermus* wasps, in which multiple females can lay eggs on a host but the females still only produce 7% males (Fig 1B; Tang *et al.* 2014; Lupi *et al.* 2017; Abdi *et al.* 2020a,b,c). These cases therefore suggest that identifying additional factors that favour female-biased sex ratio is required.

A possible explanation for the observed female-biases is that there is the potential for mutually beneficial cooperative interactions between females (Schwarz 1988; Stark 1992; Komdeur *et al.* 1997; Cronin & Schwarz 1997; Schwarz *et al.* 1998; Martins *et al.* 1999; Clutton-Brock 2002; Tang *et al.* 2014; Kapranas *et al.* 2016).For example, in presocial, allodapine bee *Exoneura bicolor*, cooperative nesting occurs among related females, which result in higher per capita reproductive outputs (Schwarz 1988; Cronin & Schwarz 1997). In this case, a more female biased offspring sex ratio can be favoured, to increase these beneficial interactions between related females, as a form of local resource enhancement (LRE; in this literature we focus on LRE provided by females). Cooperative interactions beetween females have been suggested to be important in both *Melittobia* and *Sclerodermus* wasps either (Abe *et al.* 2003, 2014; Tang *et al.* 2014; Lupi *et al.* 2017), but the extremely female-biased sex ratios under LRE in these species remain to be formally explained.

We expand existing theory to examine whether LRE can explain the extremely female biased sex ratios that have been observed in *Melittobia* and *Sclerodermus* wasps. We examine three factors that may be especially relevant to the biology of these species: (1) competitions between sons and between daughters; (2) the cooperative interactions that can occur at different times, either when laying eggs (intra-generational LRE) as in Pen & Weissing (2000) and Wild (2006), or when colonizing females help each other before competition (trans-generational LRE); (3) both females and males may disperse to different extents, hence varying the degree to which these competitive and cooperative interactions occur locally We specifically look at how sex-specific dispersal rates, the number of foundresses, and fecundity effect of LRE influence the evolution of sex ratios.

## Models and Analyses

### Life cycle

We assume Wright’s (1931) island model of dispersal, in which the metapopulation is subdivided into an infinite number of patches each fostering *n* mated females. We focus on a particular female, and we denote her proportional investment of reproductive resource into sons (“sex ratio’) by *x*_•_, the average sex ratio of the adult females in her patch in the same generation by *x*_0_, and the average sex ratio of adult females in the metapopulation by *x*. Immediately upon birth, juvenile males may disperse to an alternative patch at a rate *d*_m_ each, or else stay in the natal patch (1 − *d*_m_), followed by random mating on the patch, with each female mating only once but each male potentially mating many times. Males die after mating and females disperse with a probability of *d*_f_ each. After dispersal, breeding females compete for the limited number of breeding sites on the patch (*n*), after which the metapopulation is returned back to its original size and a new cycle starts (Fig 2A).

**Figure 2:**
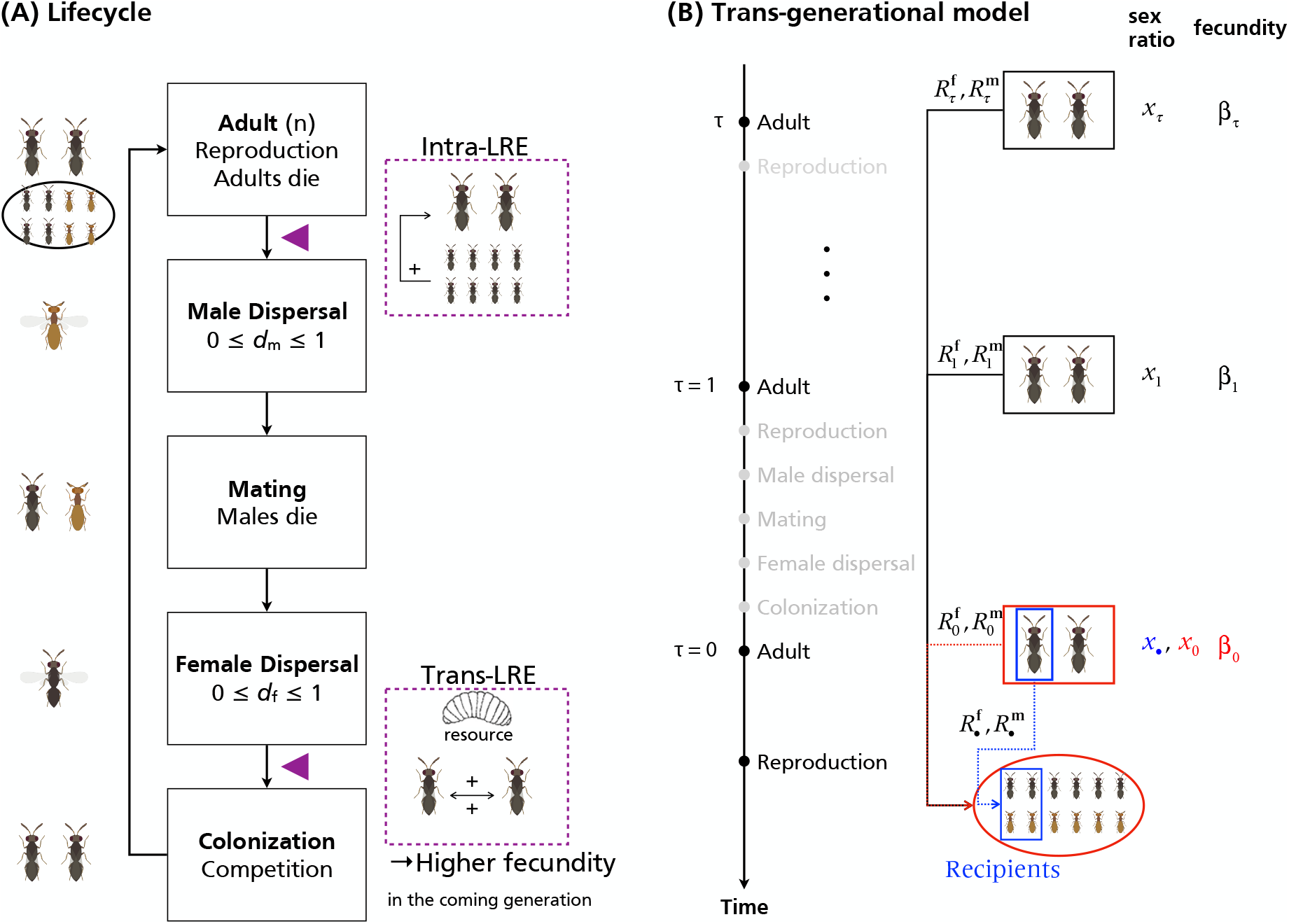
Schematic illustration of (A) lifecycle and (B) trans-generational relatedness. (A) Gray individuals: females. Brown individuals: males. Purple triangle: possible timing of LRE. Model species: *Melittobia australica* (but note that males of this species really are of flightless; Matthews *et al.* 2009). (B) Adult within blue box: the focal individual; juveniles within blue box: the focal individual’s offspring; red: average in the patch. We count the generations backwards in time (*τ* = 0 the present, *τ* = 1 parental, etc). *R* are relatedness coefficients, each from the corresponding actor’s perspective (arrows).

We consider two types of LRE. In the first, LRE occurs due to helping behaviors among juvenile females (before dispersal) that promotes the survival rate of all juveniles born in the same patch. We refer to this as “intra-generational LRE’ (“intra-” indicates that survival rate depends on the sex ratio, *x*_0_, of the focal generation, *τ* = 0, where *τ* ≥ 0 is a generic symbol to count the generations backwards ordinally in time: *τ* = 0 for the present, *τ* = 1 for the parental generation, and generally *τ* for *τ*-generations prior, and we refer to “*τ*-th generation’ henceforth). Intra-generational LRE may be relevant in species where juvenile females engage in helping behaviors before dispersal (at the latest), or equivalently, where juvenile females assist reproduction of the reproductive adults (survivorship and fecundity interpretations are mathematically equivalent). In the second model, we posit that LRE occurs due to mutual helping at the colonization stage (before competition for breeding spots), namely “trans-generational LRE’ (“trans-” indicates that the fecundity depends on the sex ratio, *x*_1_, of the previous generation, *τ* = 1). This applies to species where females communally colonize common patches, as in *Sclerodermus*.

### LRE by natal juvenile females: intra-generational LRE

We start with our analyses for intra-generational LRE, where offspring can help members of their parents. We assume that, for a patch with the average sex ratio *X*, the per capita fecundity (which is the number of offspring born times their survival rate) is given by *β*(*X*). Turning our attention to the focal patch with its inhabitants’ average sex ratio *x*_0_ (including her own sex ratio), the fecundity of the females in the patch is given by *β*_0_ ≔ *β*(*x*_0_), and the average fecundity (the total number of offspring produced per capita times their survival rate; we use ‘≔’ to define a quantity henceforth) in the metapopulation assuming that the mutants are vanishingly rare is given by *β*° ≔ *β*(*x*). Using a parameter *α* (with 0 ≤ *α* ≤ 1) which tunes the strength of LRE on fecundity (*β*_0_), we formulate *β* by:

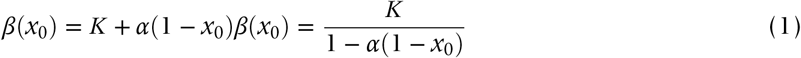

(see Appendix A for derivation), where *K* is a baseline of per capita fecundity in the absence of LRE. *β*_0_ is larger when neighboring individuals produce more females (*x*_0_ lower). The fecundity *β*(*x*_0_) decreases from *K* /(1 − *α*) to *K* as *x*_0_ varies from 0 to 1, and grows from *K* to *K*/*x*_0_ as *α* varies from 0 to 1 (for *x*_0_ > 0 fixed).

### LRE by colonizing females after dispersal: trans-generational LRE

We now turn our attention to trans-generational LRE, where juvenile females of the same generation can cooperate after dispersal for communal colonization. We use the same symbol (*β*_0_) to designate the fecundity of individuals in the focal patch, to keep the consistency with the intra-generational LRE model. We write *x*_*τ*_ for the average sex ratio of adult females in the focal patch in the *τ*-th generation, and *β*_*τ*_ for their fecundity (Fig 2B). We recursively define *β*_*τ*_ by:

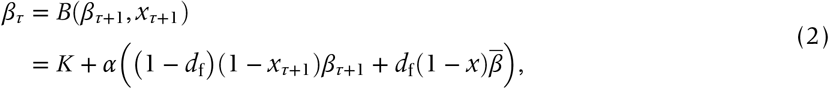

where *α* (with 0 ≤ *α* ≤ 1) measures the strength of LRE as before, 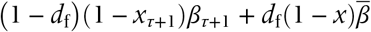 is proportional to the density of females after female dispersal (before competition), and 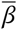 is the metapopulation-wide average of *β* to be determined by assuming that it has reached a stable equilibrium value for a phenotypically monomorphic population with *x*. The equilibrium value for 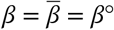 is given as the solution to *β*° = *B*(*β*°, *x*); that is:

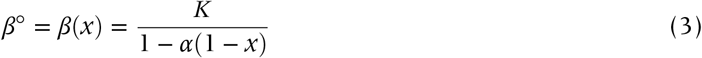

(see Eqn (1)), which is always locally stable for given *x* (i.e., *β*_*τ*_ converges to *β*° given *x* is fixed). Specifically, the fecundity of the focal female in the present generation *τ* = 0 is given by:

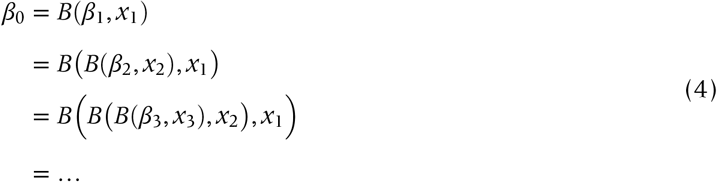

(see Appendix A for more details), which implies that to determine (the effect of selection on) *β*_0_, we need to consider an expected sequence of retrospective sex ratios in the focal patch, (*x*_1_, *x*_2_, *x*_3_, …), in addition to the focal’s and neighbors’ sex ratios in the present generation, (*x*_•_, *x*_0_) (Lehmann 2007, 2008). LRE supplied by founding females hence generates the trans-generational kin selection effects in viscous populations, by which the impacts of biased sex ratios in the patch descend down to the reproductive success of individuals (including the focal’s offspring) living in future generations, which thus in turn induces selection on the sex ratios.

### Invasion fitness and the selection gradient

We can write the invasion fitness of the focal female through daughters and sons (respectively) as:

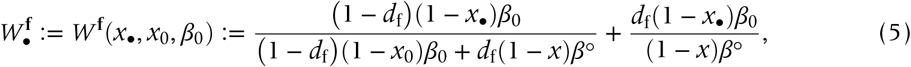

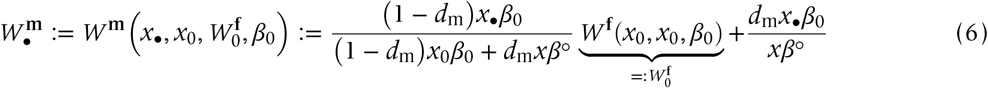

(see Lehmann 2007; Gardner *et al.* 2009; see Appendix B1–3 for derivation), where the invasion subcomponent for sons (Eqn (6)) is envisioned as a function of the focal adult female’s sex ratio *x*_•_, patch-average sex ratio *x*_0_, the survival rate of a random female as a mate for local males 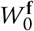 (local mating bonus; see Frank 1998, pp.199), and the average fecundity of the focal adult female *β*_0_. Note that *β*_0_ depends on model assumptions (intra- and trans-generational LRE).

We perform the neighbor-modulated fitness approach to kin selection methodology (Taylor & Frank 1996; Frank 1998; Rousset & Billiard 2000; Rousset 2004; Taylor *et al.* 2007), particularly for sex-structured populations (Taylor 1990; Taylor *et al.* 2007; Gardner *et al.* 2009) with trans-generational effects of kin selection (Lehmann 2007, 2008). We take a random juvenile female and male in the present generation each as a recipient, and adult females breeding in the *τ*-th generation (with *τ* = 0, 1, 2…, including the focal juveniles’ mother) each as an actor.

To ease biological interpretation, we here posit that *σ*_RC_ ≔ (1 − *d*_f_)^2^ tunes the intensity of LRC, which is mathematically the probability that the focal adult female’s daughters compete for resources with a juvenile female born in the same patch (eqns (7) and A20 in Wild & Taylor 2004). Similarly, the intensity of LMC is proportional to *σ*_MC_ ≔ (1 − *d*_m_)^2^, which is the probability that the focal adult female’s sons compete for mates with a juvenile male born in the same patch. Increasing *σ*_RC_ (or *σ*_MC_) favours less (or more) female-biased sex ratios (respectively). Also, the effect of extra daughters born locally on males’ reproductive success is given by *σ*_MB_ ≔ (1 − *d*_m_)(1 − *σ*_RC_) (i.e., local mating bonus; see Frank 1998, pp.199), which reads as the probability that males mate locally (1 − *d*_m_) times the probability that the mated female(s) do not encounter local resource competition with a juvenile female born in the same patch 1 – (1 − *d*_f_)^2^ = 1 − *σ*_RC_.

Using these *σ*_RC_, *σ*_MC_ and *σ*_MB_, the condition for which a slightly larger sex ratio (i.e., producing more sons than does the metapopulation average) is favoured by selection is captured by Hamilton’s rule:

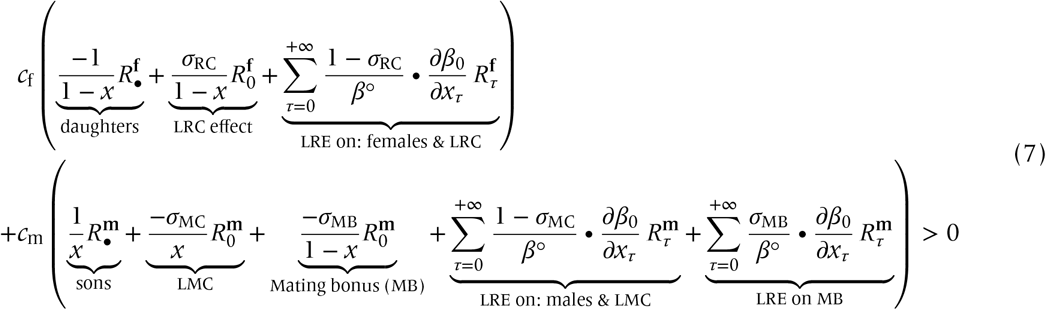

(see Appendix B4–7 for derivation; Lehmann 2007, 2008), where each derivative is evaluated at phenotypic neutrality (*x*_•_ = *x*_0_ = *x*_1_ = ··· = *x*). In Eqn (7), 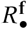 (or 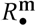) represents the regression coefficient of relatedness (‘relatedness’ hereafter; Michod & Hamilton 1980; Grafen 1985) for a juvenile female (or juvenile male) from the perspective of their mother each in the present generation (subscript *τ* = 0); 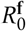 (or 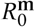) represents the relatedness for a random juvenile female (or juvenile male) from the perspective of a random adult female in the same patch each in the present generation; and 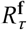 (or **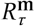**) represents the relatedness for a random juvenile female (or juvenile male) in the present from the perspective of an adult female in the *τ*-th generation (Taylor 1988; Bulmer 1994); *c*_f_ (or *c*_m_) represents the class reproductive value of female (or male; Taylor 1990; Caswell 2001).

In the absence of LRE (*α* = 0, thus ∑ terms vanishing), investing maternal reproductive resources into sons has five consequences: the decrease in daughters’ success, decrease in LRC, increase in sons’ success, increase in LMC, and decrease in MB, as in the previous studies (Taylor 1981). The summation (∑) terms each capture the sum of LRE effects each supplied by the individuals having colonized the focal patch at time epochs *τ* = 0 (for the intra-generational LRE), and *τ* = 1, 2, … (for the trans-generational LRE), on the focal female’s fitness (Fig 2B); this inclusive fitness effect occurs by which (i) LRE increases the number of sons and thus LMC, (ii) LRE increases the number of daughters and LRC, and (iii) LRE increases MB for sons.

Nullifying and solving Eqn (7) for *x* yields a candidate ESS of sex ratio (cESS henceforth; Christiansen 1991; Takada & Kigami 1991), which we generically designate with a hat 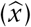.

## Results

### No LRE

We first assess the case for *α* = 0 (no LRE). By nullifying Eqn (7) for 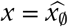 with *α* = 0 gives:

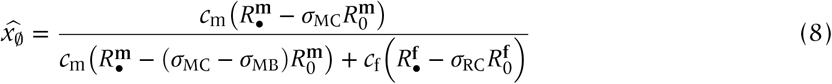

in agreement with (Wild & Taylor 2004). This expression clarifies how the local interactions affect the adaptive value of sons and daughters, where remember that *σ*_RC_ = (1 − *d*_f_)^2^, *σ*_MC_ = (1 − *d*_m_)^2^, and *σ*_MB_ = (1 − *d*_m_)(1 − *σ*_RC_). Note that for males, the total scale of competition, which includes the effect of LRC among the males’ mates (i.e., females that thus received males’ gametes), reads *σ*_MC_ − *σ*_MB_, which is negative when (1 − *d*_f_)^2^ < *d*_m_ < 1 (null for either *d*_m_ = 1 or *d*_m_ = (1 − *d*_f_)^2^; otherwise positive). Eqn (8) is a general expression of cESS under LRC and LMC (but without LRE) when male dispersal precedes mating and subsequent female dispersal (“DMD model’ in Wild & Taylor 2004). Substituting equilibrium values of the relatedness shows that 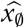 exhibits overall a female- or male-bias when *d*_m_ is small or large (respectively) and it approaches 1/2 (Fisherian sex ratio) as *n* increases (Fig 3A).

**Figure 3:**
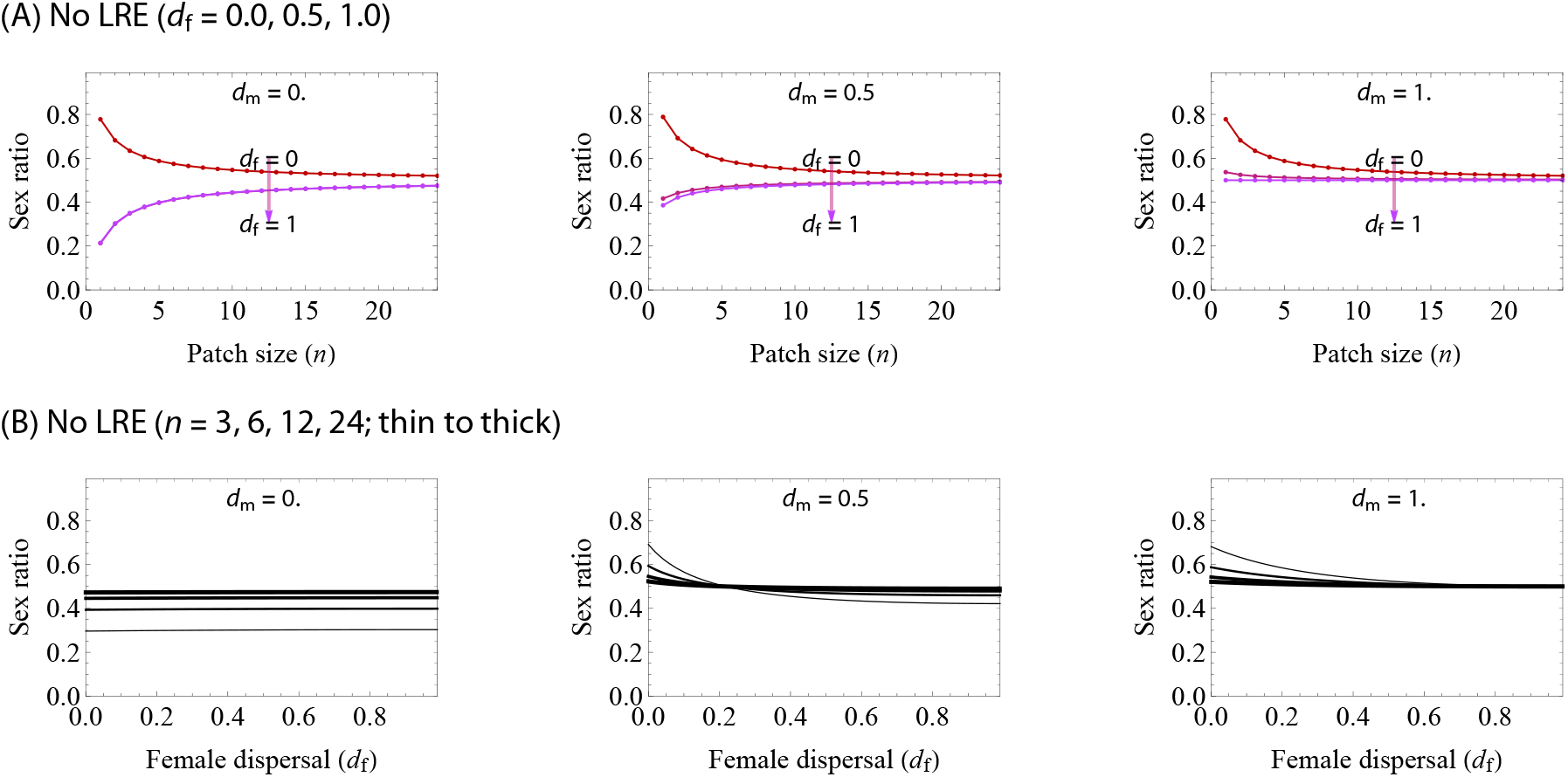
Evolutionary outcomes of sex ratio without LRE under haplodiploidy. (A) Sensitivity to *n*. Larger *n* monotonically tends to favour Fisherian sex ratio. Increasing *d*_f_ is likely to favour less male-bias. Note that *d*_f_ = *d*_m_ = 0 (red curve in the left panel) is an exceptional case in which all patches are mutually isolated, and this case therefore invalidates the present analyses (instead, entailing stochastic analyses). For (slightly) positive values for *d*_f_ > 0, the evolutionary outcome show very weak sensitivity to female dispersal rate (purple curves heavily overlapped in the left panel, which are visually difficult to separate from each other). (B) Sensitivity to *d*_f_ (with *d*_m_ = 0, 0.5, 1 from left to right panels, and *n* = 2, 4, 8, 16 from thin to thick curves). Generally, high group sizes (*n*) favour Fisherian sex ratio. When male dispersal is completely limited (left panel), sex ratio is almost invariant with *d*_f_. Increasing *d*_m_ results in a shift to male-bias when female dispersal is small, and as *d*_f_ increases, female-bias is likely to be favoured by selection (middle). When male dispersal is complete (*d*_m_ = 1), the resulting sex ratio is male-biased and approaches Fisherian (1/2) as *d*_f_ increases. All figures produced by nullifying Eqn (7).

As in the classical LMC theory, inserting *d*_m_ = 0 (no male dispersal as in *Melittobia*; Matthews *et al.* 2009) yields *σ*_MC_ = 1(> *σ*_RC_) and *σ*_MB_ = 1 − *σ*_RC_, meaning that *σ*_MC_ − *σ*_MB_ equals *σ*_RC_ on the denominator of Eqn (8); this subsequently supplies:

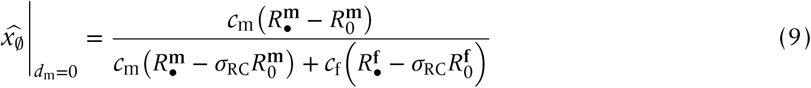

(Appendix B; eqn 3 in Gardner *et al.* 2009). Particularly for haploids and diploids, we get 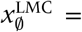 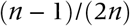 regardless of the female dispersal rate (Hamilton 1967; Bulmer 1986; Frank 1986; Taylor 1988; Bulmer 1994; Frank 1998; Gardner *et al.* 2009). This dispersal-invariance is due partly to the concomitant effects of producing more daughters on weaker LMC but stronger LRC with these effects exactly canceling one another out (Taylor’s (1992) cancelling principle; Wilson *et al.* 1992; Taylor 1992). For haplodiploids, Eqn (8) does depend on dispersal rate but in a negligibly minor manner (almost-invariance in female dispersal; Fig 3B, left panel).

In the case for *d*_m_ > 0, male biased sex ratios may occur when *d*_f_ is small, in contrast to the almost-invariance result for *d*_m_ = 0 (Fig 3B). Increasing *d*_m_ > 0 (say 0.5) leads to strongly male-biased sex ratios yet with a possibility of female-bias when *d*_f_ is relatively large (Fig 3B). When *d*_m_ = 1 (*σ*_MC_ = *σ*_MB_ = 0), only is male-bias the candidate evolutionary outcome (Fig 3B; see also SI figure in Appendix C). Overall, we find that sex ratio tends to bias towards the more dispersing sex (Bulmer & Taylor 1980; Taylor 1994; Wild & Taylor 2004).

### Effects of LRE: general patterns

We now consider the consequences of LRE (*α* > 0). We found three general patterns. First, both types of LRE drive the evolution of more female-biased sex ratio and less male-biased sex ratios (Fig 4A; see also SI Fig 1). Second, the effect of intra-generational LRE is stronger than is trans-generational LRE (Fig 4B); more precisely, the effect of intra-generational LRE is independent of sex dispersal propensities of both sexes, while that of trans-generational LRE decreases with the dispersal rates of both sexes. Finally, *d*_m_ = 0 (no male dispersal) as in classic LMC theory predicts that this special case leads to “almost-invariance results’, in which sex ratio is insensitive to female dispersal rate (Hamilton 1967; Bulmer 1986; Frank 1986; Taylor 1988; Bulmer 1994; Frank 1998; Gardner *et al.* 2009).

**Figure 4:**
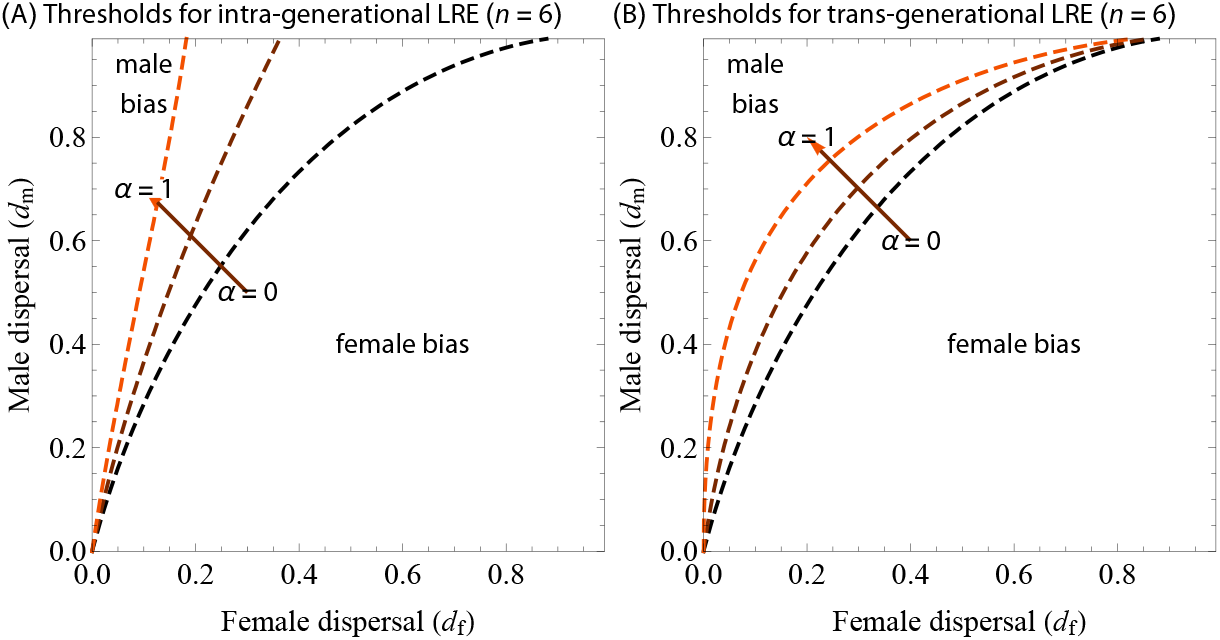
Threshold condition for the biased sex ratio at cESS. Each contour represents the condition for Fisherian sex ratio (1/2) to be cESS and separates the region for female- and male-biased sex ratios. Female biased sex ratios (bottom right zones) become more likely as *α* increases (where threshold curves plotted for *α* = 0, 0.5 and 1). The contours are produced using Eqn (7) with *x* = 1/2 (Fisherian sex ratio) inserted.

### Intra-generational LRE

We deal with general values of dispersal rates (*d*_f_ and *d*_m_, each ranging between 0 and 1), but will make an exception for *d*_m_ = 0 (no male-dispersal), because the results for *d*_m_ = 0 are qualitatively different from the results for general values 0 < *d*_m_ ≤ 1. The other advantage of presenting the specific result for *d*_m_ = 0 is that this assumption gives a simple formula, comparable with the previous theoretical work (Hamilton 1967; Bulmer 1986; Frank 1986; Taylor 1988; Bulmer 1994; Frank 1998; Gardner *et al.* 2009), and applies to many species like *Melittobia* (Matthews *et al.* 2009). As such, we present the results for *d*_m_ = 0 and 0 < *d*_m_ ≤ 1 separately; note that *d*_m_ = *d*_f_ = 0 means that patches are completely isolated from each other and entails stochastic analyses (Sigmund 2010), and so we omit this possibility.

We find that Hamilton’s rule (Eqn (7)), which assesses the direction of selection, is equal to:

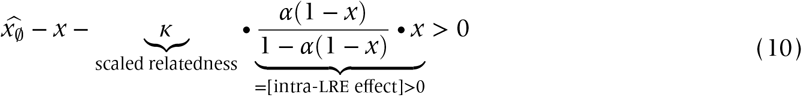

where *κ*, referred to as ‘scaled relatedness’, measures the extent to which the extra juveniles produced via LRE are likely to share the common ancestor (see Appendix B for more precise interpretation), as a function of *n*, *d*_f_, *d*_m_, in reference to the expected strength of kin competition (van Cleve 2015). The last term represents the effect of intra-generational LRE on the inclusive fitness of the focal individual. Clearly, with LRE, cESS is smaller than 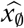, that is, LRE leads to more female-biased sex ratios (Fig 4A).

We numerically evaluated the cESS to find that larger group sizes favour less female biased sex ratio and the cESS eventually approaches 1/2 (or Fisherian sex ratio) as *n* → +∞ (Fig 5, left panels). Increasing *α* leads to more female-biased sex ratios (Figs 4 and 5). As in the results for no LRE, sex ratios may be biased towards the more dispersing sex, but LRE makes the evolution of female-biased sex ratios more likely.

**Figure 5:**
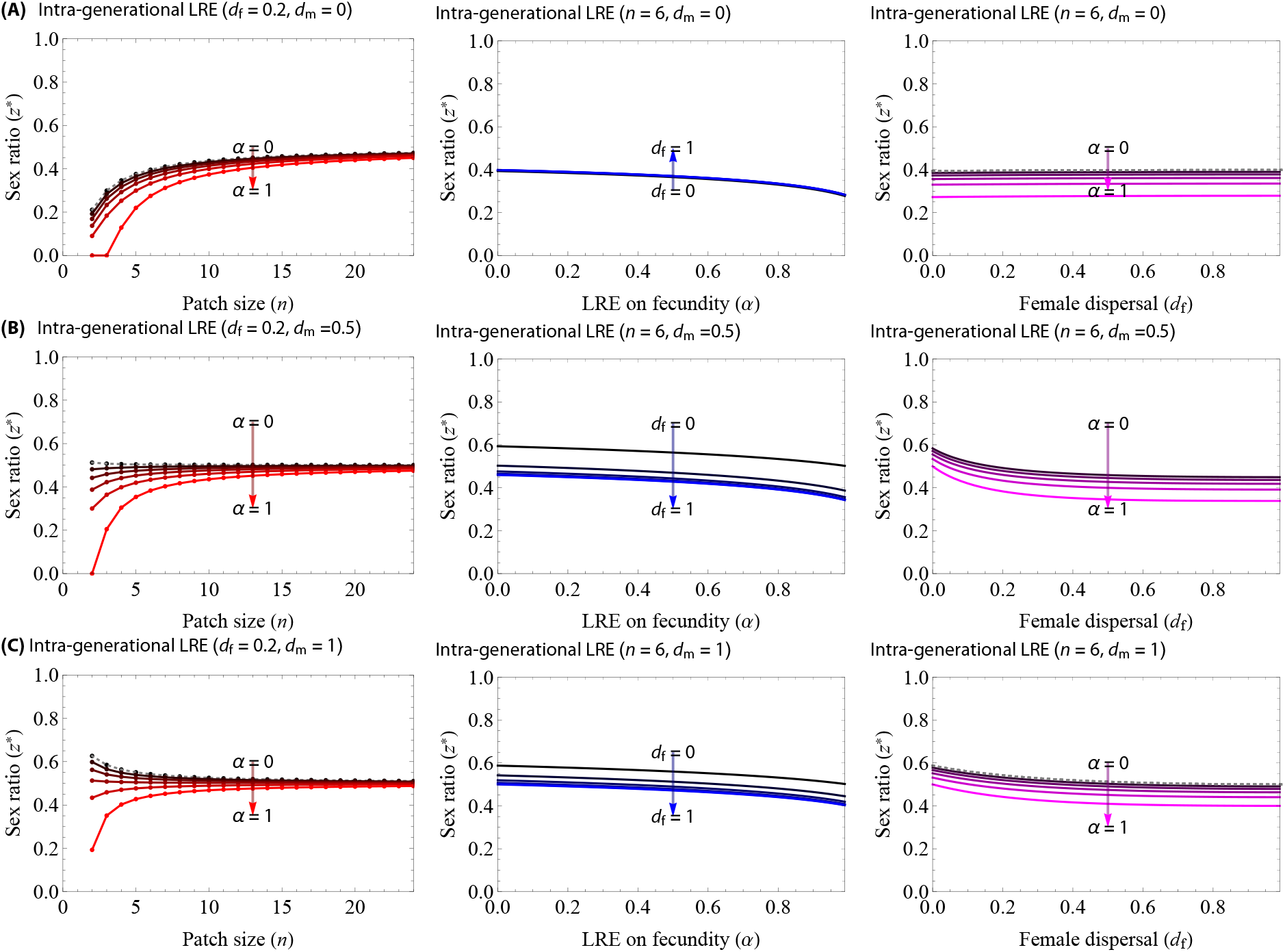
cESS under *d*_m_ = 0, for intra- generational LRE (top panels, A) and trans-generational LRE (bottom panels, B), under haplodiploidy. Gray dots: results under no LRE.

#### Example: no male-dispersal, *d*_m_ = 0

Suppose for now *d*_m_ = 0, and in this case, we can show that *κ* = 1/*n* (Taylor 1992; Gardner *et al.* 2009) and therefore ploidy has no influence on the effect of LRE nor *κ* (Taylor 1992; Lehmann 2007). This is partly because males are fully philopatric, mating takes place in prior to female dispersal, and female dispersal allows males’ and females’ gametes both to disperse by the same degree, which leads to *σ*_MC_ − *σ*_MB_ = 1 − *d*_f_ ^2^ ≡ *σ*_RC_ (i.e., males and females are subject to the same degree of local competition), where ‘≡’ is identity (“always equivalent to’). This scenario is similar to plants undergoing gametic (pollen) and zygotic (seed) dispersal when pollen dispersal is fully restricted within a patch (see Rousset 2004; Ravigné *et al.* 2006; Iritani 2020 for more details). That is, decomposing the scale of competition tells us otherwise missed fact: when *d*_m_ = 0, the scale of competition for both sexes is equivalent (thus giving the almost-invariance result).

#### Varying male-dispersal, *d*_m_ > 0

Now we tune *d*_m_ from 0 to 1 and assess its impacts upon cESS. We find that increasing *d*_m_ or *d*_f_ is likely to favour less or more female-biased sex ratios (Fig 5; respectively), and taking both to 1 leads to Fisherian sex ratio. For an intermediate male dispersal (*d*_m_ = 0.5), male-bias is still likely but with a possibility of switching from male- to female-bias as *α* or *d*_f_ increases. Therefore, under intra-generational LRE, the sex ratios, which could be otherwise male biased, may be biased towards female by natural selection.

### Trans-generational LRE

We find that trans-generational LRE also facilitates the evolution of female-biased sex ratio (Fig 4B). Numerical estimation revealed that larger group sizes favour less female biased sex ratio and the cESS eventually approaches 1/2 (or Fisherian sex ratio) as *n →* +∞ (Fig 6, left panels), and increasing *α* leads to more female-biased sex ratio (Fig 6), as in the trans-generational LRE model. The inclusive fitness effect of trans-generational LRE decreases with *d*_f_; when *d*_f_ = 1 (full female dispersal), for instance, the effect of trans-generational LRE vanishes for any *α* > 0 (Fig 6A, midel panel). The trans-generational LRE, which ensues after female dispersal, is sensitive to *d*_f_ because the probability that females can help their relatives (by remaining in the natal patch, 1 − *d*_f_) decreases with *d*_f_.

**Figure 6:**
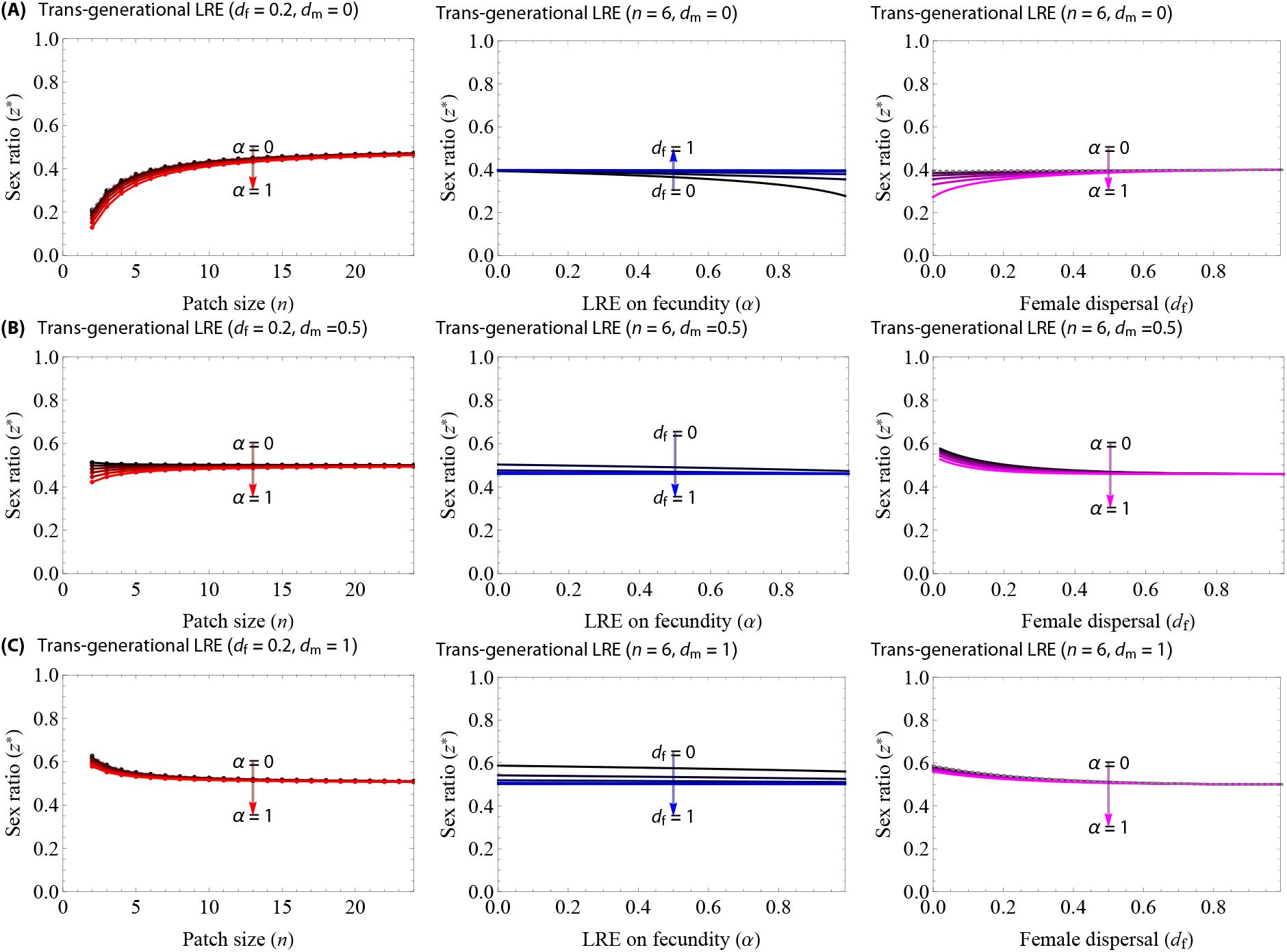
cESS under *d*_m_ = 0, for intra- generational LRE (top panels, A) and trans-generational LRE (bottom panels, B), under haplodiploidy. Gray dots: results under no LRE.

#### Example: No male-dispersal

When *d*_m_ = 0, we find that Hamilton’s rule reads:

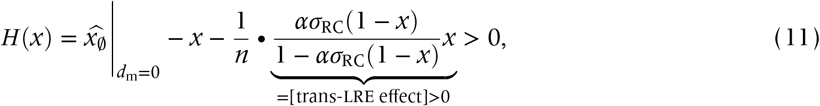

where the scaled relatedness is now given by *κ* = 1/*n* as in the intra-generational model, but the last term on Eqn (11) is clearly smaller than that on Eqn (10); i.e., trans-generational LRE is weaker than is intra-generational LRE in terms of their effects on selection.

#### Varying male-dispersal

Varying *d*_m_ > 0 turns out to give complicated form of Hamilton’s rule (see Appendix B), except for the extreme case *d*_m_ = 1:

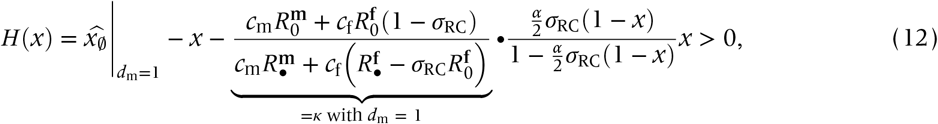

which tells us that *ασ*_RC_ in Eqn (11) is now replaced with *ασ*_MC_/2, with 1/2 meaning that only half of females’ genes are transmitted to females (who, as opposed to males all dispersing, are likely philopatric and thus potentially contribute to the build up of trans-generational relatedness). From the expression, the scaled relatedness *κ* is now with reference to null (no LMC or MB as all males disperse) for males and 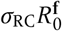 (LRC, which is zero when all females disperse *d*_f_ = 1).

## Discussion

We found that cooperative interactions between females (LRE) can lead to more female-biased sex ratios than those predicted by local mate competition (LMC) theory. Specifically, w e have considered two types of LRE, and found that cooperation from offspring to their parents generation (intra-generational LRE) can lead to more female biased sex ratios than cooperation between members of the same generation (inter-generational LRE). This difference is because intra-generational LRE occurs prior to female dispersal and therefore each juvenile female has direct access to helping genetically related juveniles, thereby increasing the inclusive fitness of producing daughters over sons; in contrast, trans-generational LRE occurs after female dispersal, and dispersed females are unable to provide help to relatives, thereby reducing the inclusive fitness benefit of LRE as *d*_f_ increases. Our key result is therefore that LRE, alongside LMC, has the potential to generate extremely female-biased sex ratios, of the form that have been observed in nature, in species such as *Melittobia* and *Sclerodermus* wasps (Abe *et al.* 2003, 2014; Tang *et al.* 2014).

As found by previous theory, we found that, in the absence of LRE, selection in general favours the biased sex ratio towards the more dispersing sex (Bulmer & Taylor 1980; Taylor 1994; Wild & Taylor 2004), which is because kin competition between members of one sex reduces the relative value of producing that sex. In contrast, when females interact cooperatively (LRE), this increases the relative value of producing females, and natural selection thus favours less male-biased or more female-biased sex ratios (Figs 4 and 5; Taylor 1981; Emlen *et al.* 1986; Pen & Weissing 2000; Wild & Taylor 2004; Wild 2006; Wild & West 2009; but see Khwaja *et al.* 2017).

Despite the formal similarity between intra- and trans-generational LRE models, there is the quantitative difference in the consequences of dispersal rates for sex ratios. For intra-generational LRE, increasing the intensity of LRE (*α*) leads to more female-biased sex ratio by increasing the benefit of producing juvenile females who assist their mother (Figs 5 and 6). This selective force acts even when female dispersal rate is high, because intra-generational LRE allows juvenile females to assist their own mother before dispersal. In contrast, the trans-generational LRE predicts that increasing the intensity of LRE (*α*) has a weaker effect on the selection for the female-bias (Figs 4 and 6) compared to the intra-generaional LRE. The inclusive fitness effect of trans-generational LRE vanishes if females undergo complete dispersal; in other words, *d*_f_ = 1 implies that cESS is independent of *α* for the trans-generational LRE. This result is because following complete female-dispersal, dispersed juvenile females (the proportion *d*_f_) do not have the access to their relatives and there are unable to engage in helping. Hence, the two models suggest that distinguishing the timing of LRE is of crucial importance for sex ratio evolution in empirical and experimental systems.

Our models provide a formal theoretical explanation for the extreme sex ratio biases that have been observed in *Sclerodermus harmand*, and several *Melittobia* wasp species. In both these cases, local mate competition is likely, but the offspring sex ratios are much more female biased than would be expected from local mate competition theory (LMC; Fig 1). We have shown in our trans-generational model that a combination of cooperative interactions between sisters (LRE) and LMC can lead to more female biased sex ratios, consistent with the empirical data. In *Sclerodermus harmandi*, females cooperate to suppress hosts and engage in brood cares (Tang *et al.* 2014; Kapranas *et al.* 2016; Lupi *et al.* 2017). In *Melittobia*, females tend to aggregate on the same hosts and cooperatively parasitise them (J. Abe, unpublished; Rosenheim 1990), females fight against the symbiont mites of host species (Okabe & Makino 2008), and their female offspring collaborate to tunnel into the materials of host nests to disperse (Deyrup *et al.* 2005), suggesting that LRE may promote female-biased sex ratios. However, in *Melittobia australica*, the quantitative discrepancy appears to be large (Figs 1 and 6). Possible factors may include multiplicativity of LRE *β*(*x*) = *K*/(1 − *α*(1 − *x*)^*θ*^) (as in Appendix C2 and SI Fig 3; reader may want to compare this formula with Eqns (1) and (3)) which lead to even more female-biased sex ratios and LRE from juvenile females that specifically increases the production of daughters (not sons), or alternatively, we may need to consider cohesive dispersal or kin recognition among females that promotes LRE but reduces LRC. Recent field work shows that sex ratios in *Melittoiba* depend on female dispersal status, in which sex ratios produced by dispersing females are less female-biased than ones by non-dispersing females or observed in the laboratory (Fig. 1), but more female-biased than the prediction by a model incorporating dispersal status (Abe et al. in preparation). It may even be the case that *Melittobia australica* are unable to facultatively adjust sex allocation in response to local patch size due to some constraints (Shuker & West 2004; Greeff *et al.* 2020). Despite this, our models offer a useful framework to study the evolution of sex ratios under LRE and allow further development of theoretical work.

Our models are also applicable to a wide range of other organisms, in which there can be both cooperative (positive) and competitive (negative) interactions with relatives. For example: (i) mammals, birds and insects where offspring help their parents, but can also compete for breeding sites (territories; Komdeur *et al.* 1997; Clutton-Brock 2002; Stark 1992); (ii) allodapine bees where sisters cooperate to form nests, but offspring can compete for resources (Schwarz 1988; Cronin & Schwarz 1997; Schwarz *et al.* 1998); (iii) *Diadasina distincta* bees where individuals must compete for nest sites, but nesting at higher densities can reduce parasitism (Martins *et al.* 1999). In all cases, our general prediction is that a female-biased sex ratio bias is favoured to increase cooperative interactions and reduce competitive interactions.

To conclude, our analysis suggest that intra- and trans-generational LRE from females does promote female-biased sex ratios, but their impacts upon the evolutionary outcomes differ especially in terms of the consequences of dispersal: intra-generational LRE does not depend on female dispersal rate but trans-genearational LRE does. One of the possible extensions of the present model is to study joint evolution of sex ratio and other traits under LRE (Mullon *et al.* 2016, 2018). For instance, how does joint evolution shape the association between sex-biased dispersal and sex allocation strategy, e.g., in birds and vertebrates (Frank 1990; Komdeur *et al.* 1997; Goltsman *et al.* 2005; Banks *et al.* 2008; Hjernquist *et al.* 2009)? Future studies could be directed towards more realistic modeling of LRE, by e.g., incorporating the effects of the number of adult females (*n*) on LRE, or working on intra-sexual LRE (‘who helps whom’; Rodrigues & Gardner 2013; Rodrigues & Kokko 2016). Working with specific organisms of interest based upon multiple approaches may yield a better understanding of the evolution of sex ratios in viscous populations.

## Acknowledgement

We thank Troy Day, Ian Hardy, Yoshitaka Kamimura, Asher Leeks, Joyce Tong, and anonymous reviewers for helpful comments on various versions of the manuscript; Ian Hardy, Hiroaki Yanagisawa, and Baoping Li for sharing their data; JSPS-KAKENHI (grant numbers 19K22457, 19K23768, and 20K15882 to RI), and ERC (grant number 834164 to SW) for funding. We thank Laurent Lehmann for kindly hosting RI in 2015 and his group for discussion, which helped RI develop the mathematical analyses presented here.

## Author contribution

JA conceived the idea; RI carried out the mathematical analyses; JA and RI drafted the first version of the manuscript; all authors contributed to the revision.

## Appendix

## A Fecundity mediated by LRE

## A-1 Derivation

Let us for now focus on the intra-generational model. Pick a focal patch and denote the average phenotype on that patch by *x*_0_; on that patch, the fecundity per capita may be the sum of (i) baseline fecundity (denoted by *K*), and (ii) effects of LRE; the latter occurs from helping behaviors of daughters, and the total number of females per patch is given by *n* • (1 − *x*_0_) • *β*(*x*_0_), in which such an effect is equally shared by *n* females and thus should be divided by *n* (therefore *n* being canceled), which hence yields:

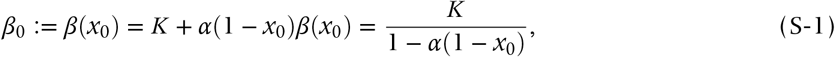

as in the main text (Eqn 1).

The same logic gives the corresponding recursion for the trans-generational model, by modifying the equation above to:

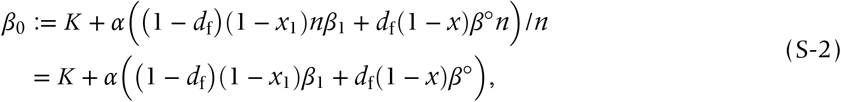

which may be reasoned as follows: the first term is a baseline fecundity; the second term overall represents the effect of LRE; the fraction 1 − *d*_f_ of females derives from the natal patch, in a density (1 − *x*_1_)*β*_1_*n*; *d*_f_ from other patches, in a density (1 − *x*)*β*°*n*. Assuming that such LRE effect is shared among *n* individuals, we divide this term by *n*, obtaining the above expression. We can see that *β*_0_ is a function of (*β*_1_, *x*_1_), and *β*_1_ is a function of (*β*_2_, *x*_2_) (by the same reasoning), and so forth; this means that we need to deal with the nested function of *β*_0_, based upon Lehmann (2007, 2008). With this nestedness, we henceforth write:

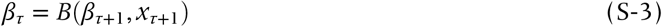

for *τ* ≥ 0.

When *x*_*τ*_ = *x*_•_ = *x* (for any *τ* ≥ 0; neutrality condition), we have

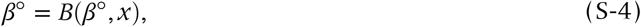

which immediately gives:

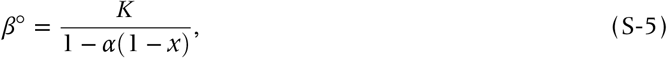

whereupon we recover Eqn (3) in the main text.

To guarantee *β*° be locally stable, we separate the timescales (holding *x*_•_ = *x*_*τ*_ = *x* for all *τ* ≥ 0) by assuming the sequence of the average fecundity values 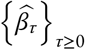 (with a hat put to emphasize quasi-equilibrium) is, using Eqn (S-2), determined by:

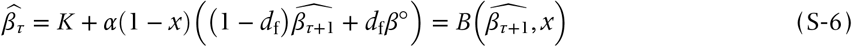

from which we have:

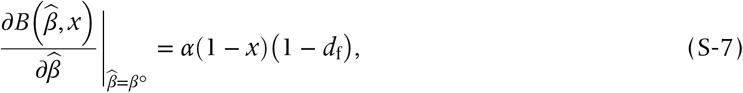

which is obviously positive and smaller than 1, thereby guaranteeing the local stability (NB: there may be a case in which this quantity is unity but only when *x* = *d*_f_ = 0 and *α* = 1; the necessary condition *x* = 0 is excluded and thereby we focus on strict inequality).

## A-2 Effects of selection

## Intra-generational LRE

Using Eqn (S-1),

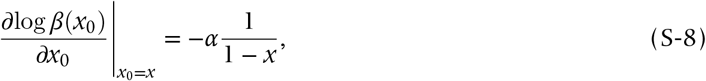

and thus:

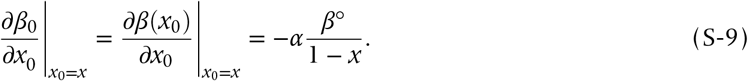

## Trans-generational LRE

From Eqn (S-2), similar computation gives:

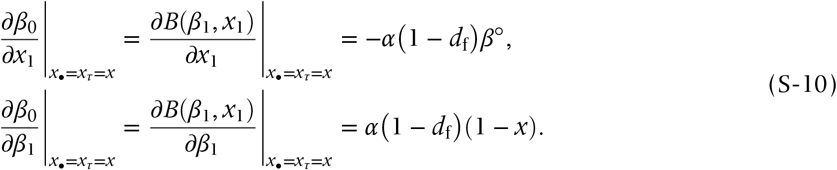

Induction yields:

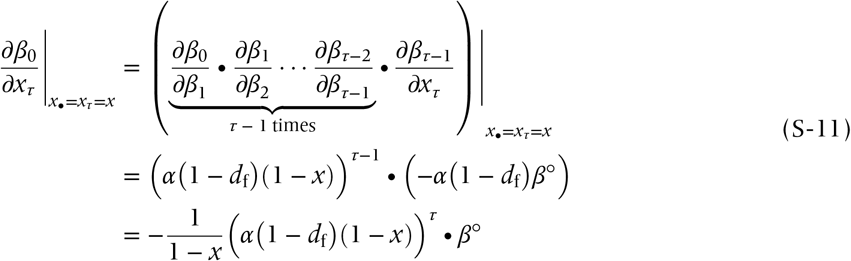

for *τ* ≥ 1. That is, the effect of selection through the expected average of sex ratios in the focal group of the *τ*-th generation decays geometrically in *τ*.

## B. Invasion fitness and the selection gradient: general

The reproductive success of a focal individual occurs through her daughters (winning the breeding opportunity) and her sons (winning the mating opportunity), and therefore we separate these two terms explicitly by using the class reproductive values (Taylor 1990; Caswell 2001). We use the subscripts in and tr to distinguish the two models.

## B-1 Success via daughters

First of all, the focal female can produce the total number *β*_0_ of eggs. A proportion 1 − *x*_•_ of those eggs develops into female. Therefore we have (1 − *x*_•_)*β*_0_ of daughters born to the focal. Pick one of the daughters;

- With a probability of 1 − *d*_f_, she stays in her natal patch, ending up with competing with, on average, (1 − *d*_f_)(1 − *x*_0_)*β*_0_ of juvenile females born in the same patch, plus *d*_f_(1 − *x*)*β*° of juvenile females each born in a different patch.
- With a probability of *d*_f_, she disperses to an alternative patch to compete with (1 − *x*)*β*° of juvenile females.

Therefore, we have:

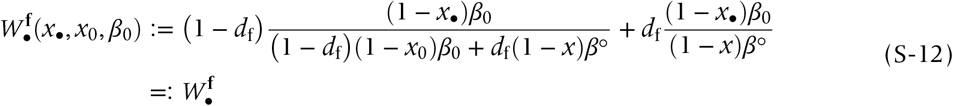

as displayed in Eqn (5) of the main text. At phenotypic neutrality (i.e., *x*_•_ = *x*_0_ = *x* and *β*_0_ = *β*°), we have 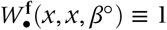 (where, by ≡, we mean “always equal to” or identity).

## B-2 Success via sons

The same logic gives success via sons. The focal female produces *x*_•_*β*_0_ of sons who may compete against on average (1 − *d*_m_)*x*_0_*β*_0_ + *d*_m_*xβ*° of juvenile males by staying philopatric (which occurs with a probability of 1 − *d*_m_); in which case, whether the male gamete survives to the next generation or not now relies on whether the mated female wins a breeding opportunity or not. Because mating is fully random within the patch, 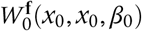 gives the proportional success of the sons’ mating partners. If the sons instead disperse, which occurs with a probability of *d*_m_, then they compete with on average *xβ*° of males, and their mating partners’ success is 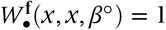. Therefore,

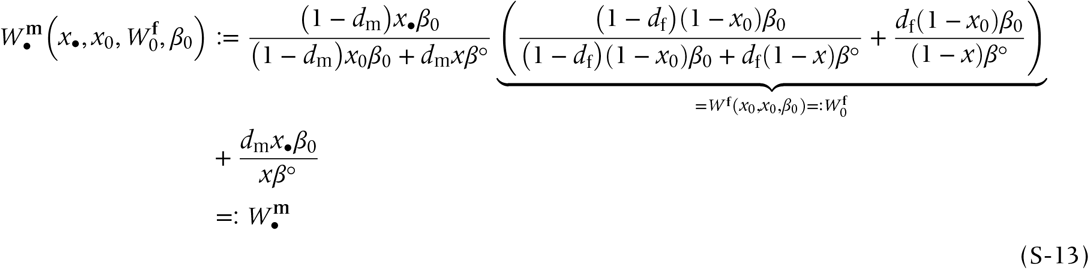

as displayed in Eqn (6) of the main text.

## B-3 Total fitness

The total number of genes of the focal individual transmitted to the next generation is therefore given by 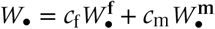 (Taylor 1990; Taylor *et al.* 2007; Gardner *et al.* 2009).

## B-4 Selection gradient (Hamilton’s rule): preliminary

Pick the locus Ξ that encodes the sex ratio and write *ξ* for the genic value of an allele of the locus from a juvenile in the focal patch, with *ξ*^**f**^ for female and *ξ*^**m**^ for male, respectively. In addition, we denote the breeding value of (i) the juvenile’s mother by 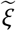, (ii) the average breeding value of the adult females in the same patch in the same generation by *η*_0_, (iii) the average breeding value of all adult females in the metapopulation by *ζ*, and (iv) the average breeding value of the adult females in the same patch *τ*-generations prior to the present by *η*_*τ*_ for *τ* ≥ 1.

Under vanishingly small genetic variation in the metapopulation, the direction of selection can be assessed by the sign of:

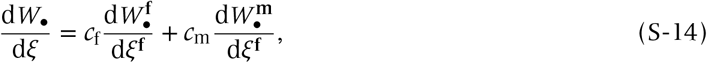

evaluated at *ξ* = *ξ*^**f**^ = *ξ*^**m**^ = *ζ*, in which *c*_f_ (or *c*_m_) is interpreted as the probability that the focal gene lineage is found in a female (or male, respectively). If the above derivative is positive, then selection favours a slightly larger allocation to males.

Further decomposition of the partial derivatives are as follows (Lehmann 2007, 2008; Gardner *et al.* 2009):

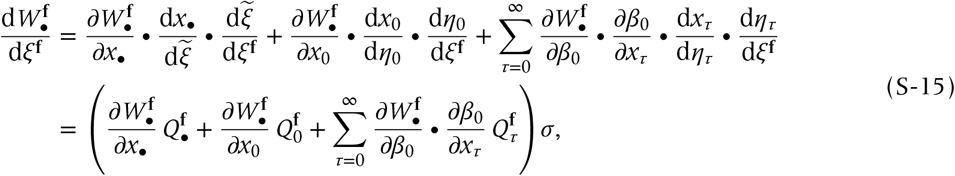

where, assuming vanishing genetic variation, we have rewritten (i) 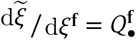 for the consanguinity of the focal female to her own daughters, (ii) 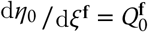 for the consanguinity of the focal female to the juvenile females born to the focal patch in the present generation, (iii) 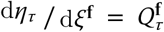 for the consanguinity of a random juvenile female and an adult female breeding in the focal patch *τ*-generations prior to the present (for *τ* ≥ 1), and (iv) *σ* is an arbitrary constant representing the slope of phenotype on genotype, given by 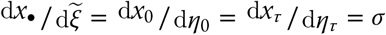, which we hereafter rescale *σ* = 1.

The similar expansion yields:

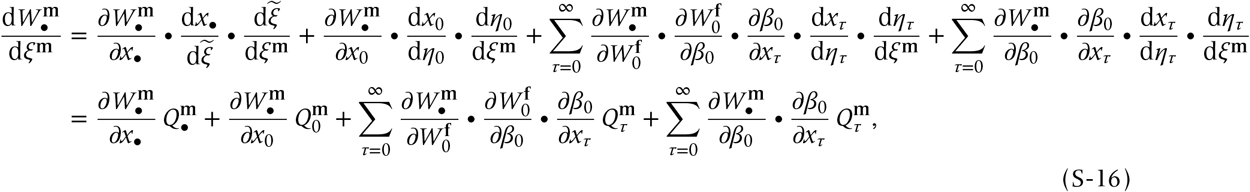

with basically the same notation as above (^**f**^ being replaced with ^**m**^).

From these, the selection gradient (that separates the intra- and trans-generational LRE effects) reads:

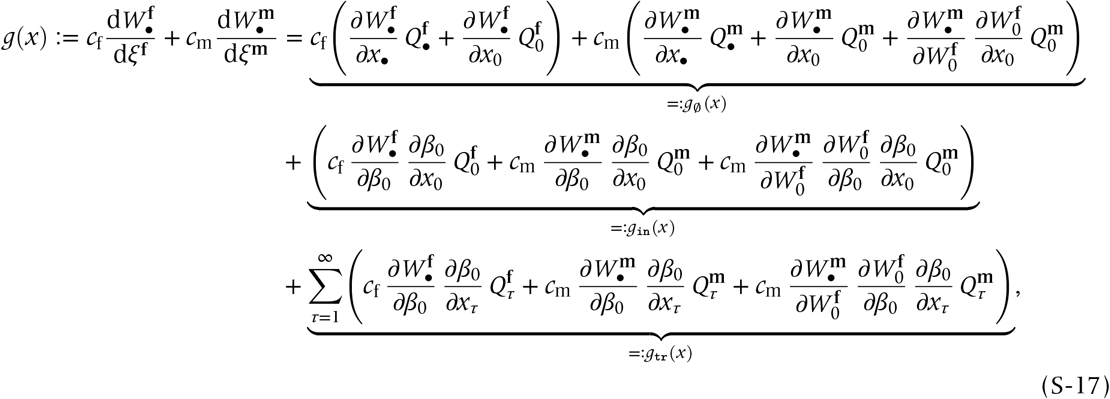

where *g*_in_(*x*) and *g*_tr_(*x*) respectively represent the subcomponent of the total selection gradient for intra- and trans-generational LRE, and *g*_∅_(*x*) represents the subcomponent of the selection gradient in the absence of LRE.

## B-5 Partial derivatives

Using Eqns (S-12) and (S-13), some algebraic manipulations give the following partial derivatives (each evaluated at neutrality):

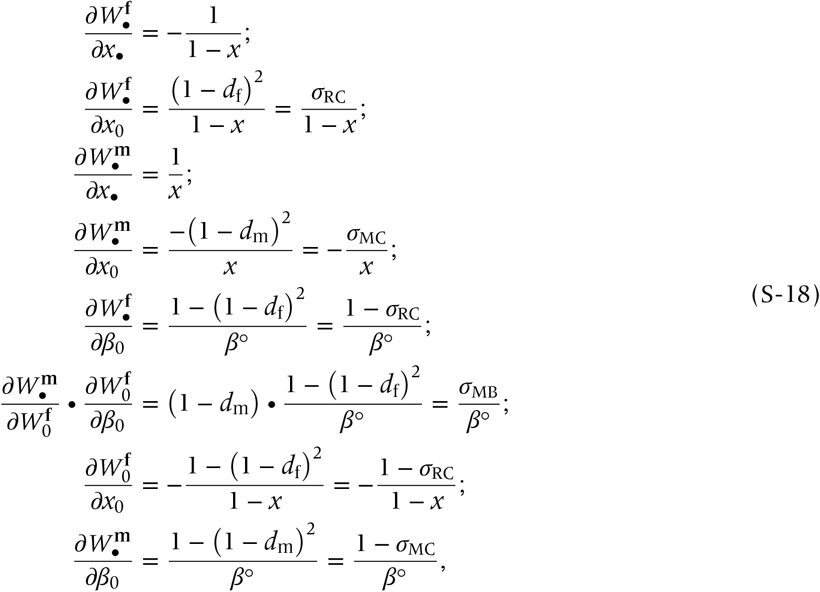

in which we have written (i) (1 − *d*_f_)^2^ = *σ*_RC_ for the strength of local resource competition among females, (ii) (1 − *d*_m_)^2^ = *σ*_MC_ for the strength of local mate competition among males, and (iii) *σ*_MB_ = (1 − *d*_m_)(1 − (1 − *d*_f_)^2^) for the strength of local mating bonus. Mathematically, these are: (i) the probability that a juvenile female competes for resource with another juvenile female born in the same patch, (ii) the probability that a juvenile male competes for mating opportunity with another juvenile male born in the same patch, and (iii) the joint probability that a male mate with a juvenile female born in the same patch and then she does escapes from competition with a juvenile female born in the same patch.

## B-6 Hamilton’s rule

If we divide both sides of Eqn (S-17) by *Q*_•_ > 0, we get:

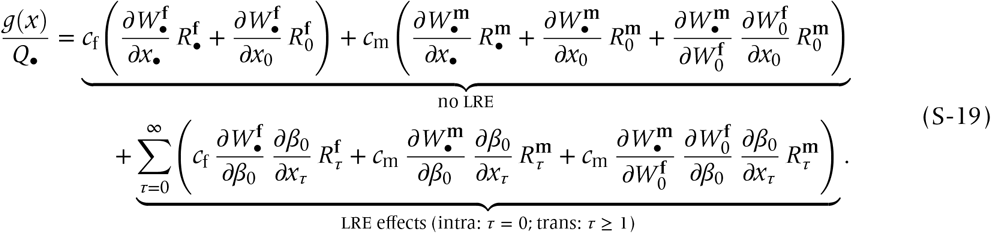

Inserting Eqn (S-18) into Eqn (S-19) without making explicit the relatedness coefficients (as functions of dispersal rates and *n*), we get:

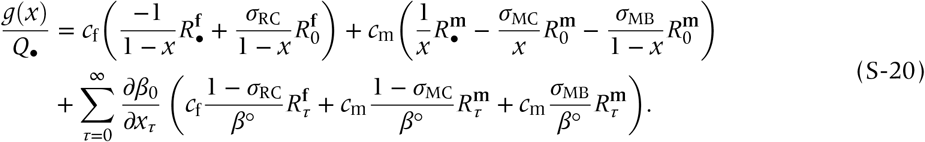

## B-7 No LRE

When *α* = 0 (no LRE), the second line of Eqn (S-20) is null. Then Eqn (S-20) vanishes if:

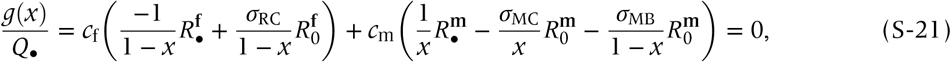

which is equivalent to:

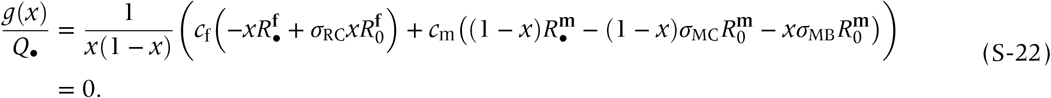

Multiplying *x*(1 − *x*) > 0 yields:

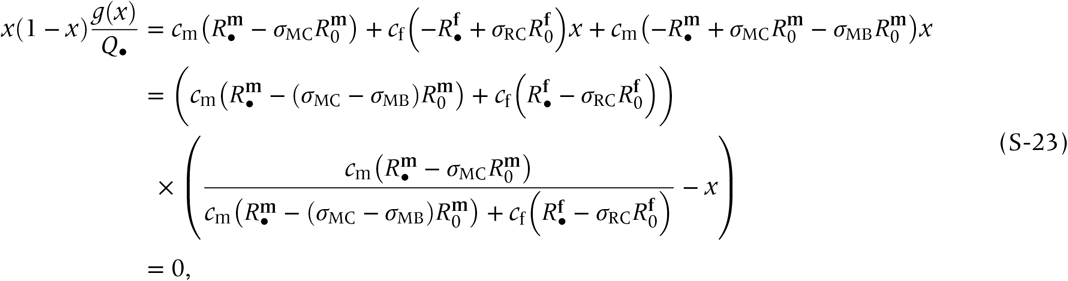

which obtains the expression for cESS (candidate ESS), of

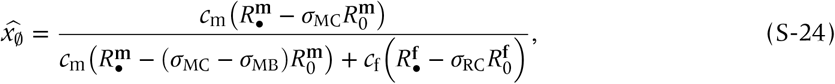

which recovers the cESS displayed in Eqns (8) and (9) of the main text. By making the relatedness
coefficients explicit with *d*_f_, *d*_m_ and *n*, we can obtain the numerical value for 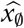 to reproduce Fig 3 of the main text.

From this manipulation from Eqns (S-21) to (S-23) we can see that it is of great use to scale *g*(*x*) by:

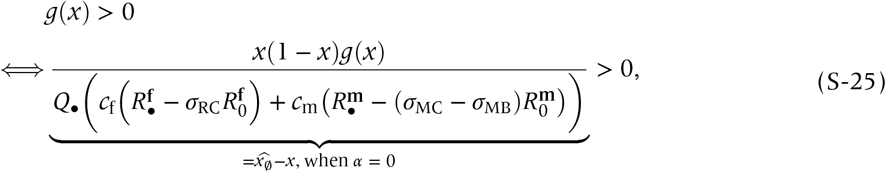

as it gives a simple measure of the direction of selection. We therefore define:

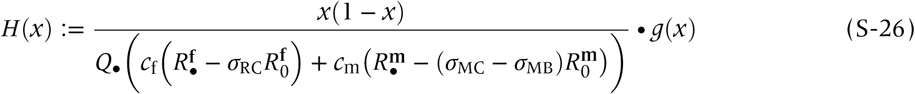

and refer to this as “scaled Hamilton’s rule,’ presented in Results of the main text (Eqns 9 – 12).

## B-8 Scaled Hamilton’s rule for the intra-generational LRE

Substituting Eqns (S-10) and (S-11) into Eqn (S-20) and rescaling it to scaled Hamilton’s rule *H*(*x*) defined in Eqn (S-26) (but without making the relatedness coefficients explicit (while extracting *τ* = 0 term on ∑) in Eqn (S-20)), we arrive at:

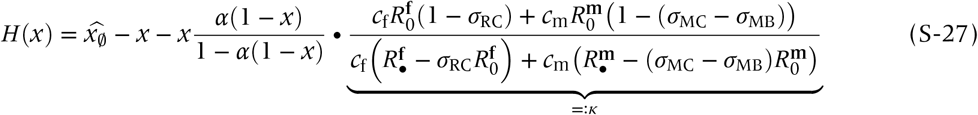

and define *κ* as above.

## B-9 Scaled Hamilton’s rule for the trans-generational LRE

Similar computation yields:

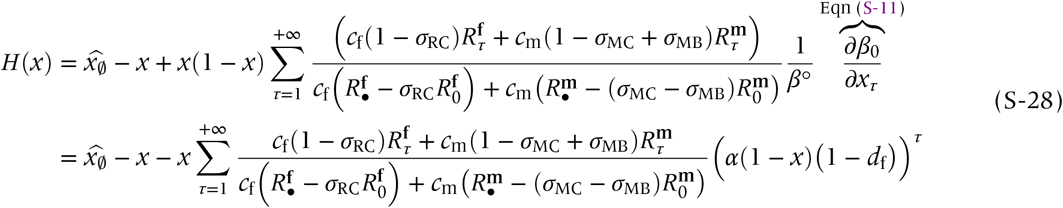

(Lehmann 2007, 2008). So we must obtain recursions for 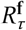, 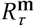 (below).

## B-10 Intra-generational consanguinities

We denote the consanguinity between (i) a juvenile female and male sharing the same patch by **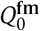**, (ii) two juvenile males sharing the same patch by 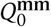, and (iii) two juvenile females sharing the same patch by 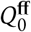, each in the same generation *τ* = 0.

Using the standard coalescent argument for haplodioloids, we get:

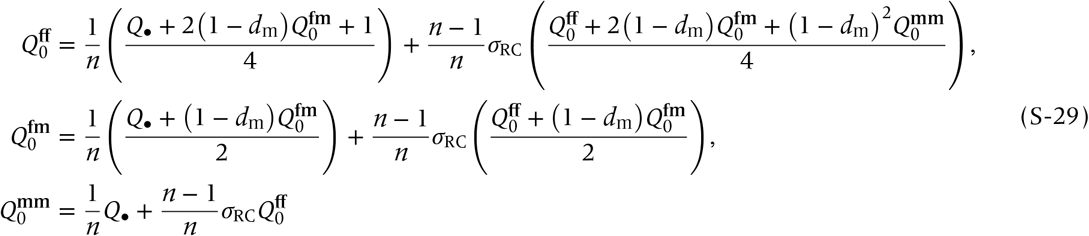

(Taylor 1988; Johnstone *et al.* 2012). Solving these to find the unique solution, we can determine each consanguinity coefficient.

The consanguinity of an adult female to

i. herself is 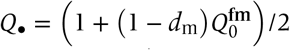,
ii. her daughter is 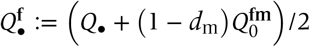,
iii. her son is 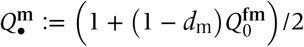,
iv. a random juveni female born in the same patch in the present generation is 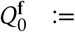 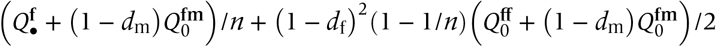, which turns out to equal 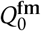, and
v. a random juvenile male in the same patch born in the present generation is 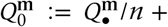 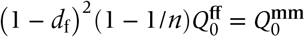.

Also, the relatedness coefficient, from the adult female’s perspective (Taylor 1988; Bulmer 1994), to

i. her daughter is 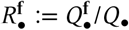,
ii. a random juvenile female on the same patch born in the present generation is 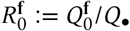,
iii. her son is 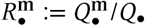, and
iv. a random juvenile male on the same patch born in the present generation is 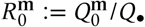.

## Proof for *κ* = 1/*n* for *d*_m_ = 0, Taylor’s cancelling principle

We here show that *d*_m_ = 0 gives *κ* = 1/*n*, which is a part of Taylor’s (1992) cancelling principle, as shown in Eqn (11).

First, plugging *d*_m_ = 0 into Eqn (S-29), we get:

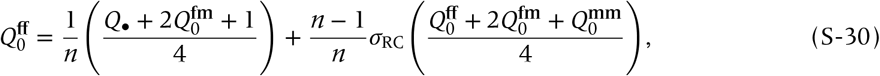

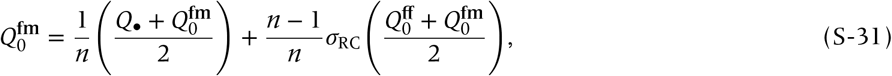

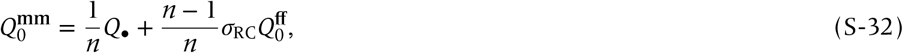

with:

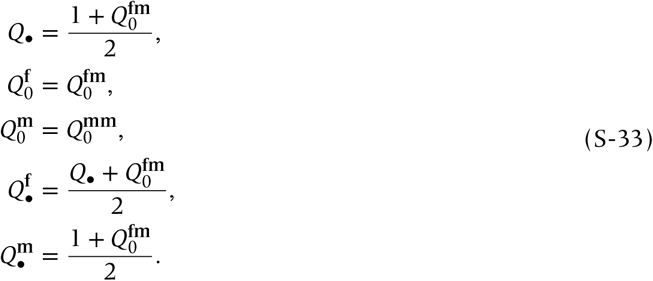

Second, computing 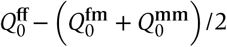 from Eqns (S-30) to (S-32) (separating LHS and RHS, while using Eqn (S-33)) yields:

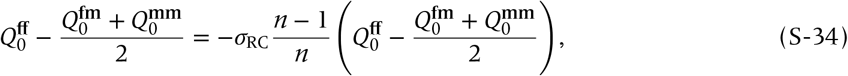

thus implying

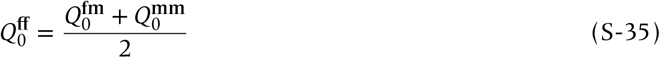

at equilibrium.

Third, let us repeat the definition of *κ* (for any *d*_m_):

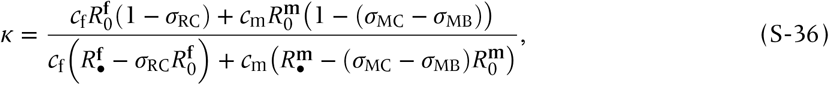

into which we substitute *d*_m_ = 0 to get:

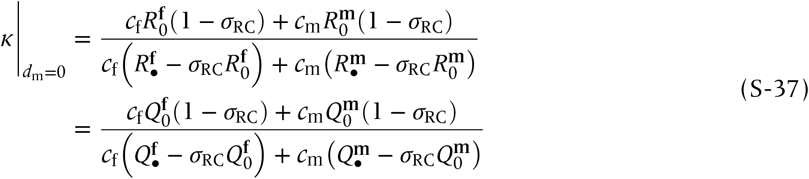

(where we used *σ*_MC_ − *σ*_MB_ ≡ *σ*_RC_ when *d*_m_ = 0). If we, for now, define 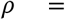 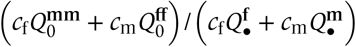, then:

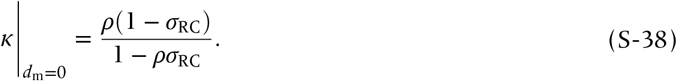

*c*_f_×Eqn (S-31) plus *c*_m_×Eqn (S-32) gives:

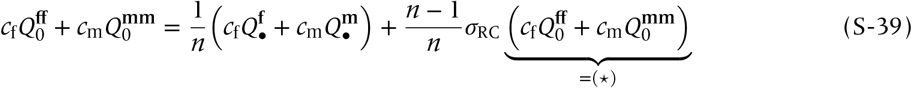

(where (⋆) is equal to 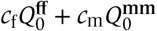 after algebraic manipulations), which, using *ρ*, implies:

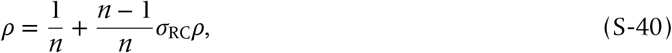

which by solving for 1/*n* gives:

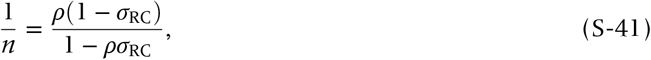

which is equal to Eqn (S-38), therefore *κ* = 1/*n* when *d*_m_ = 0. This gives the result for *d*_m_ = 0 of the main text (including Eqn 11).

## B-11 Trans-generational consanguinities

We focus on haplodiploid genetics, and designate 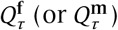 for the probability of consanguinity between (i) a juvenile female (or male) randomly sampled right after its birth in the present generation in a patch and (ii) a random adult female in the same patch in the *τ*-th generation (*τ* ≥ 0). To develop the recursion between 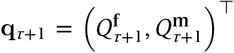 and 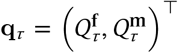 (with ^⊤^ for transp o se), we sample three classes of individuals living in the same patch but potentially in different time epochs: (i) adult females in the *τ* + 1-st generation (representatively named 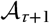), (ii) juvenile females or males born from the adult females living in the generation *τ* = 1 (named 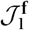 for female and 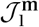 for male), and (iii) juvenile females or males born from the adult females in the current generation *τ* = 0 (named 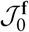 for female and 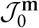 for male). The consanguinity between (i) and (ii) is by definition **q**_*τ*_ and that between (i) and (iii) is **q**_*τ*+1_. The initial condition (*τ* = 0) reads 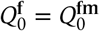 (which is the consanguinity of a random juvenile female born in *τ* = 0 and a random adult reproducing in the same generation) and 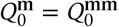 (which is the consanguinity of a random juvenile male born in *τ* = 0 and a random adult reproducing in the same generation). We depict a conceptual illustration in SI Fig 2, in which actors’ and recipients’ perspectives (or inclusive-fitness and neighbor-modulated fitness approaches) do agree, as in Taylor *et al.* (2007).

## Recursion for 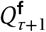

First consider the probability that 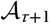 (who is a random adult female sampled from a patch in the *τ* + 1-st generation) and 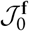 (who is a random juvenile female born from an adult female in the same patch in the present generation) share an allele of identity-by-descent (IBD). For this to occur, it entails that 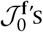 mother be of philopatric origin (1 − *d*_f_); given this, 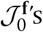 allele derives maternally with a probability of 1/2, in which case IBD probability is 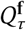; otherwise, it derives from a father (1/2) born in the same patch (1 − *d*_m_), in which case IBD probability is 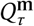. Hence,

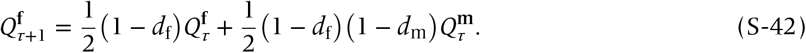

## Recursion for 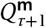

Similarly consider the probability that 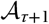 (a random adult female sampled from a patch in the *τ* + 1-st generation) and 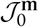 (a random juvenile male born to the adult females in the same patch in the present generation) share an allele of IBD. Given that 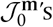 mother be of philopatric origin (which occurs with a probability of 1 − *d*_f_), 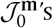 allele derives certainly maternally (probability 1), in which case IBD probability is 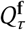, yielding:

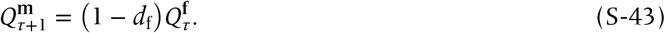

## Vector form of the recursions

In a vector form, Eqn (S-42) and Eqn (S-43), divided by *Q*_•_ read:

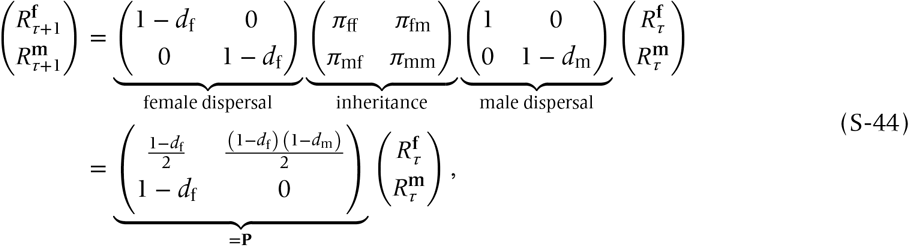

where *π*_XY_ represents the probability that a juvenile of sex X derives its gene from an adult of sex Y. *π*s are determined by ploidy levels: *π*s are all 1/2 for haploids and diploids, while *π*_ff_ = *π*_fm_ = 1/2, *π*_mf_ = 1 and *π*_mm_ = 0 for haplodiploids.

In Eqn (S-44), the first matrix determines the decay of genetic relatedness due to female emigration, in which mated females disperse and therefore males’ gametes may emigrate together (the bottom-right element should not be 1); the second describes the genetic inheritance, or movement of genes between sexes due to mating; the third represents the male dispersal, by which juvenile females’ gametes do not disperse.

Now let **V** ≔ *α* (1 − *d*_f_)(1 − *x*)**P**; then its spectral radius is lower than unity in modulus and therefore 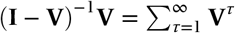 does exist (with **I** the identity). Then, Eqn (S-28) becomes:

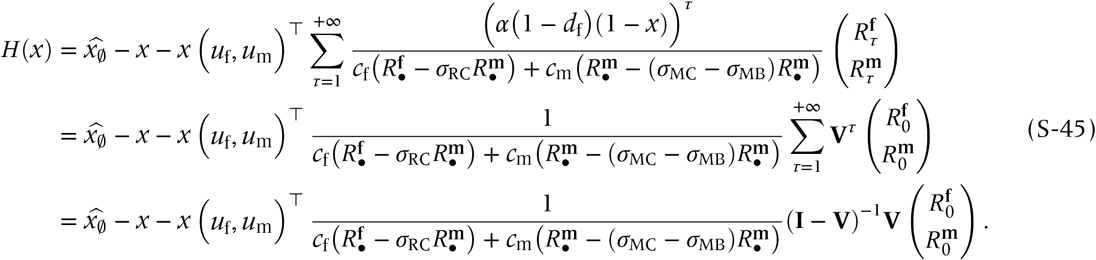

As any nonsingular bidimensional square matrix has a formula for its inverse matrix (one may want to use *Mathematica*^®^ notebook file we submitted), we can find that the trans-generational subcomponent, *g*_tr_(*x*), is given by

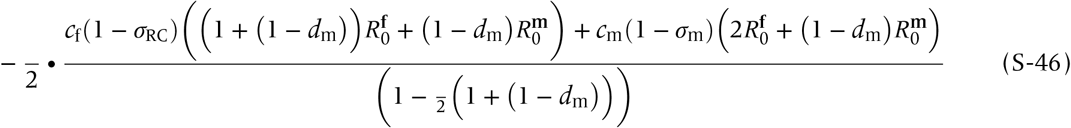

for haplodiploids, where *σ*_m_ ≔ (1 − *d*_m_)^2^ − (1 − *d*_m_)(1 − (1 − *d*_f_)^2^ and ≔ *σ*_RC_*α*(1 − *x*).

## C. Robustness under *d*_m_ = 0

## C-1 Different functional form: additive

## Intra-generational model

Suppose the fecundity of individuals in the present generation is given by:

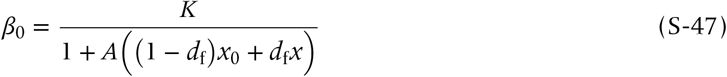

where *A* represents the strength of LRE, and we therefore have:

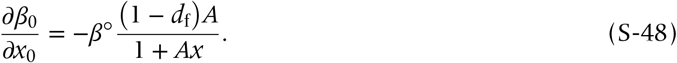

Replacing *α* in the main text with *A* = *α* / (1 − *α*) gives the corresponding analyses; therefore, the present extension is analogous to the analysis given in the main text.

## Trans-generational model

Suppose that *β* obeys the following recursion:

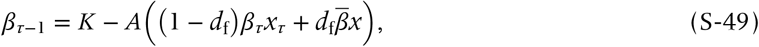

with:

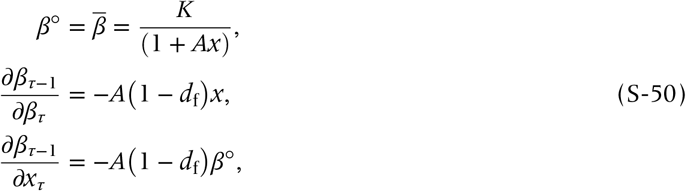

which constitutes a negative niche-construction (fecundities of the present and previous generations are negatively correlated), or aggressiveness towards offspring (of any sex) from males. In this case, a straightforward calculation similar to the analyses performed above yields:

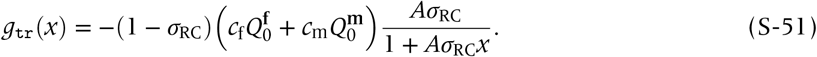

Rewriting *A* = *α* / (1 − *ασ*_RC_), we have:

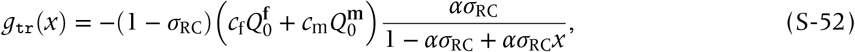

which is equivalent to the one obtained in the original, trans-generational model, with a difference in feasibility conditions: the original *α* varies from 0 to 1 while *A* may vary from 0 to 1 / (1 − *σ*_RC_) > 1, suggesting that the upper bound for the strength of LRE depends on the dispersal rate. Therefore, our results are robust against this simple recursion for *β*_*τ*_, and the original approach presented in the main text may be suitable for experimental confirmation.

## C-2 Different functional form: multiplicative

## Intra-generational model

Here we assume that LRE is generated by a multiplicative functional form given by:

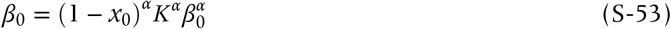

(for 0 ≤ *α* < 1), which satisfies:

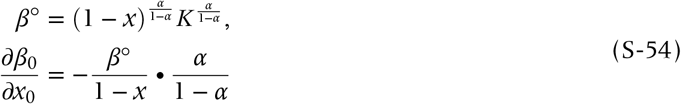

at neutrality. Therefore,

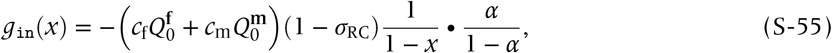

telling us that the effect of LRE is independent of the dispersal rate. Hamilton’s rule is thus given by:

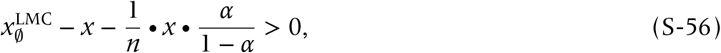

which determines the cESS as:

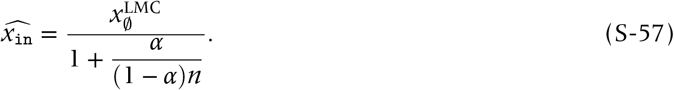

This expression immediately tells us that *α* → 1 leads to 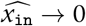.

## Trans-generational model

Similarly, consider:

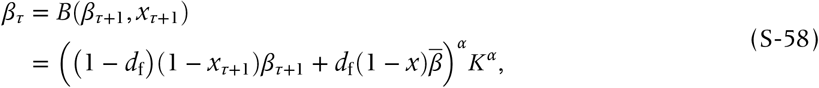

which, at equilibrium, should satisfy:

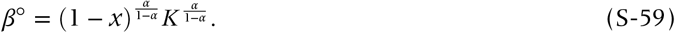

Partial differentiation gives:

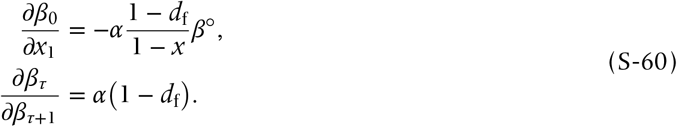

Hence,

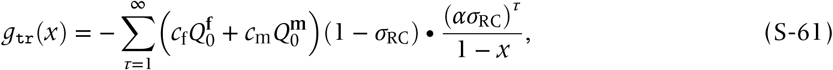

which, for 0 ≤ *α* < 1, converges to:

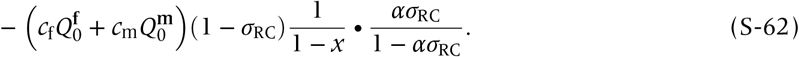

Therefore, Hamilton’s rule is given by:

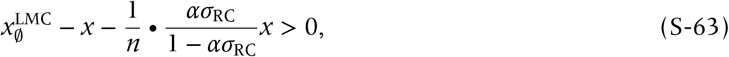

supplying the cESS, of:

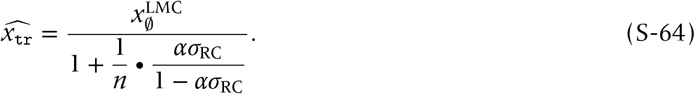

Overall, we can observe that the effects of LRE on 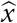 are much more pronounced compared to the additive one (SI Fig 3), which is because with this multiplicative formula *x* = 1 gives *β* = 0: producing females is a prerequisite for producing offspring. Though much simpler in any associated expressions, the model may overestimate the LRE effect. For this reason, we presented the additive formula for LRE in the main text.

## C-3 LRE from males: additive

We here demonstrate a minimal analysis for LRE supplied by juvenile males.

## Intra-generational model

Suppose that the fecundity of the females in the focal patch (with its average phenotype *x*_0_) is given by:

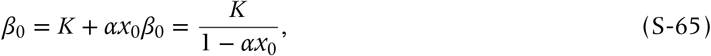

which means that:

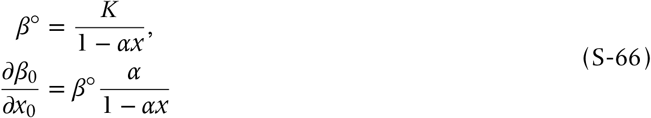

at neutrality. The preceding logic shows that Hamilton’s rule is given by:

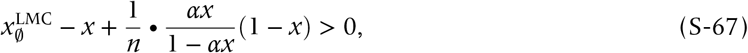

which gives:

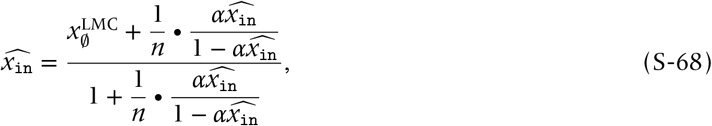

from which, by the intermediate value theorem, we can find a unique 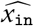 lying between 0 and 1.

## Trans-generational model

We use the function of the form:

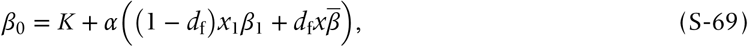

which has the equilibrium solution given by:

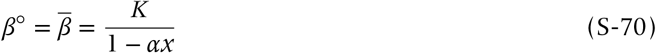

at neutrality, and satisfies:

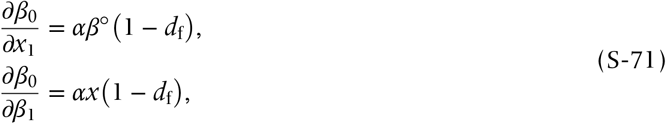

once again at neutrality. Therefore we can obtain Hamilton’s rule of:

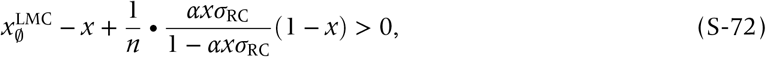

which gives the cESS 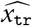 as 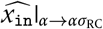 in Eqn (S-68).

## C-4 LRE from males: multiplicative

## Intra-generational model

Suppose the fecundity function of the form:

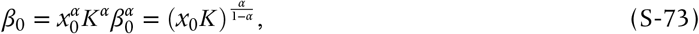

with:

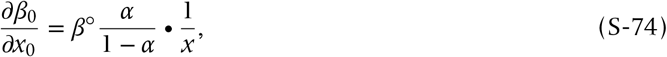

implying that Hamilton’s rule reads:

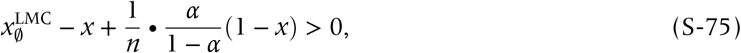

which gives:

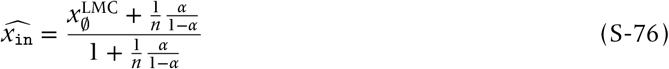

## Trans-generational model

The same logic supplies Hamilton’s rule as:

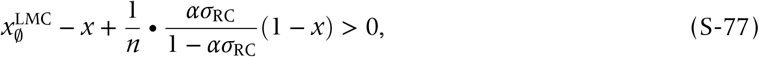

with the resulting cESS of:

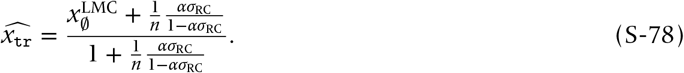

We here remark that the trans-generational LRE from juvenile males is unlikely, because males do not disperse and therefore have no access to post-dispersal, founding females. One may want to apply an alternative model to specific organisms with a specific lifecycle.

## Combining additivity and multiplicativity: Combinational LRE

We here consider

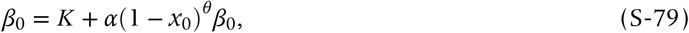

or

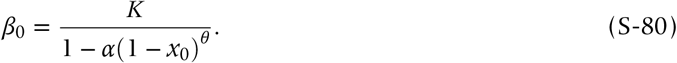

for 0 < *θ* < 1, and the resulting cESS does not have analytical formula. The numerical solutions are given in SI Fig 6.

Inserting *d*_m_ = 0 or *d*_m_ = 1 into the above equation and then rescaling the whole equation (**??**) leads to the expression given in the main text.

---

**SI Figure S1:**
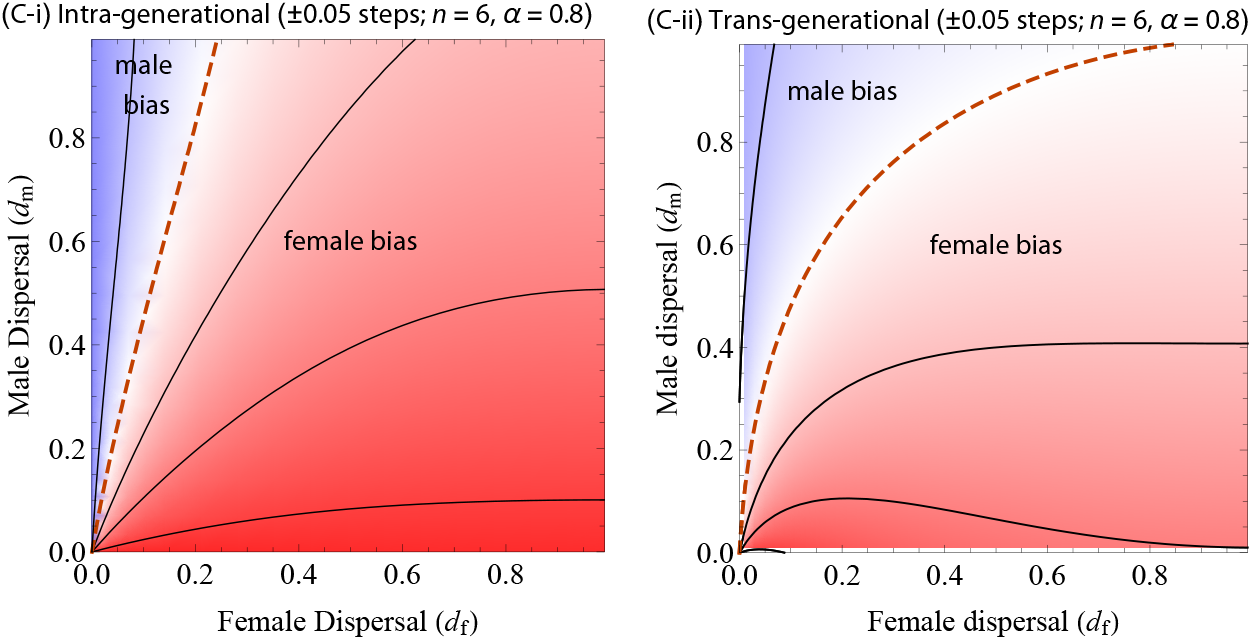
Evolutionary outcomes plotted against sex-dependent dispersal rates. Orange dotted contour: Fisherian sex ratio (*x* = .5), and the others are ±0.05 steps deviation from Fisherian.

**SI Figure S2:**
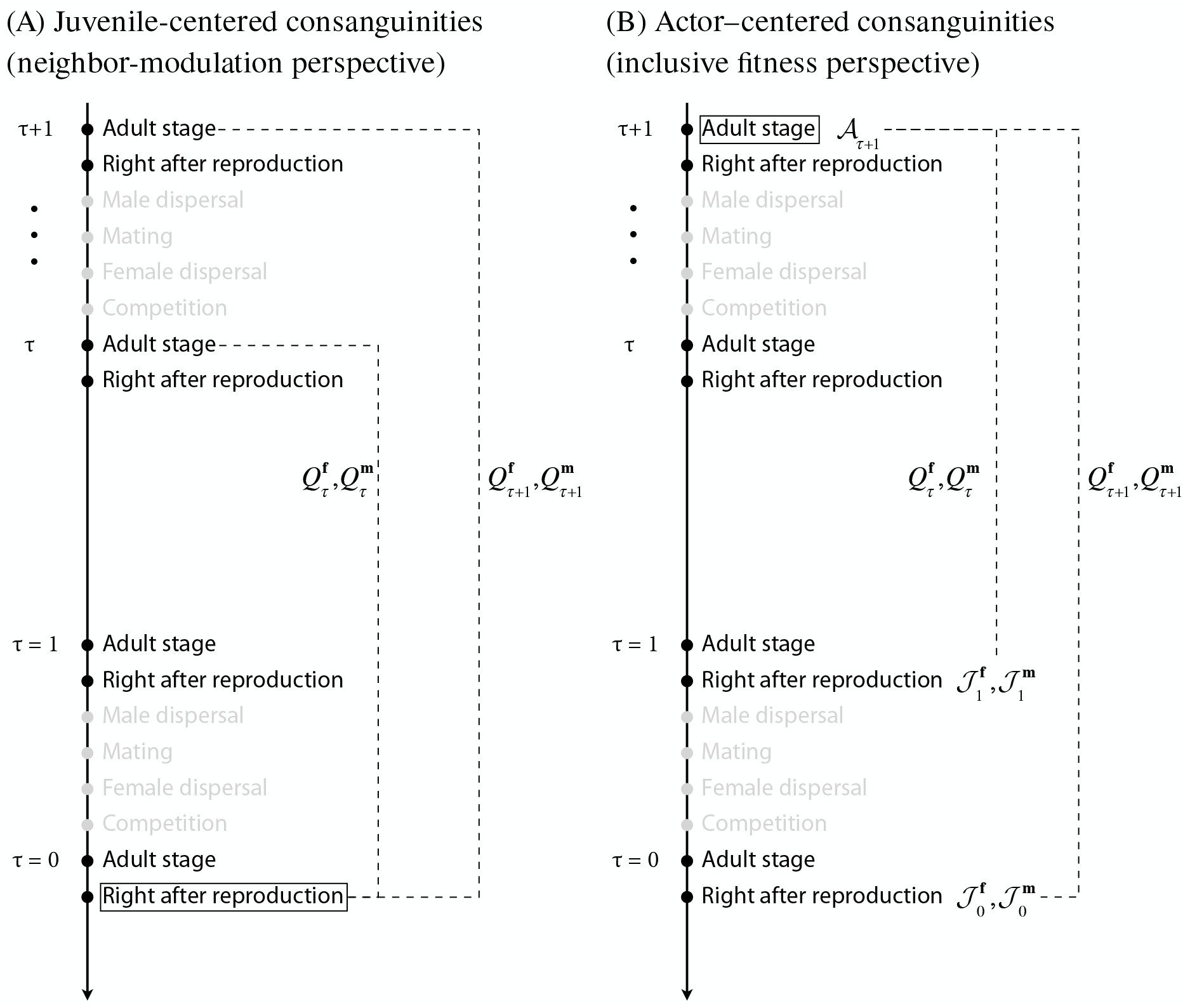
Schematic illustration to develop recursive equations for the trans-generational consanguinity

**SI Figure S3:**
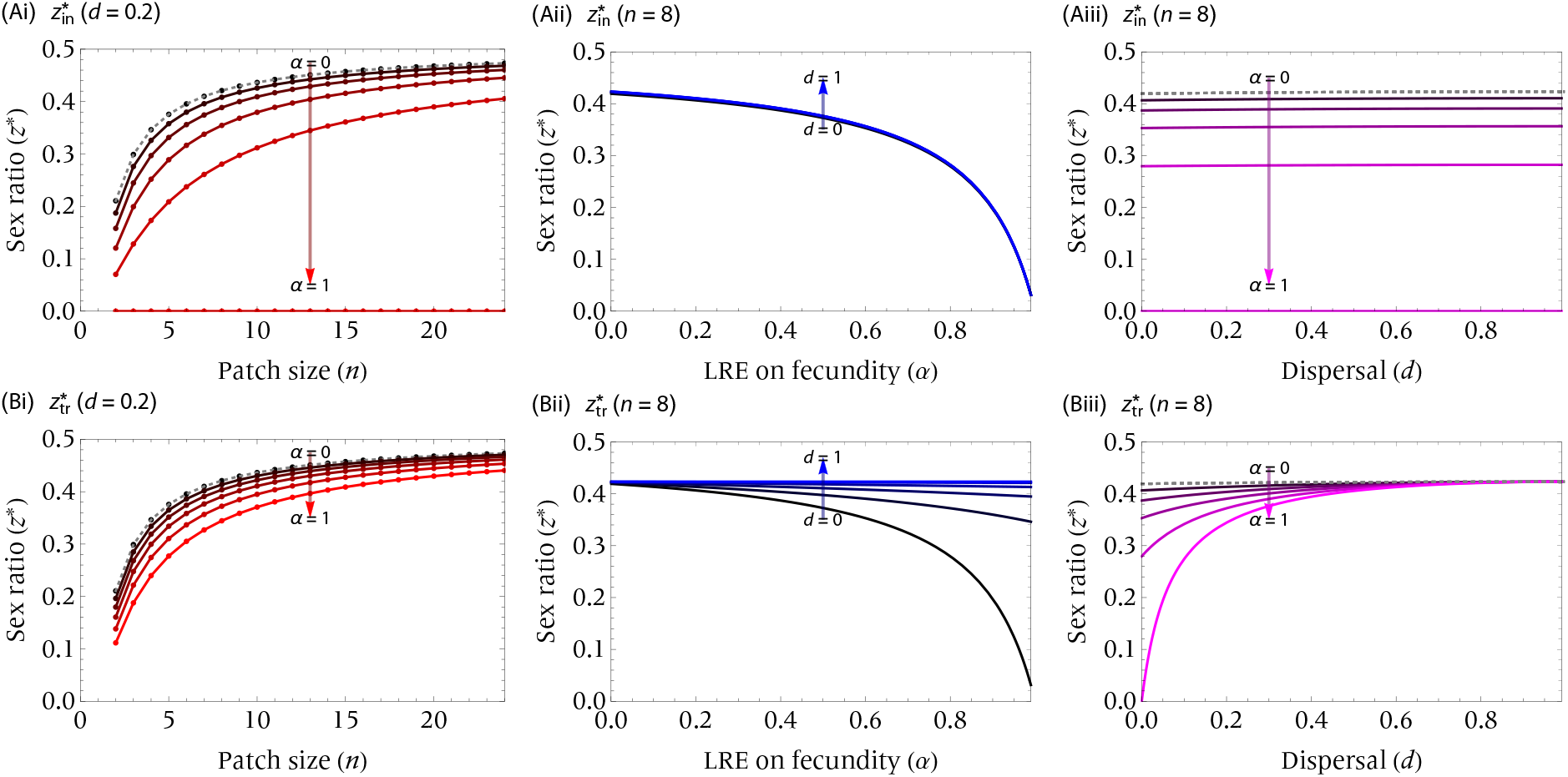
Multiplicative female-LRE.

**SI Figure S4:**
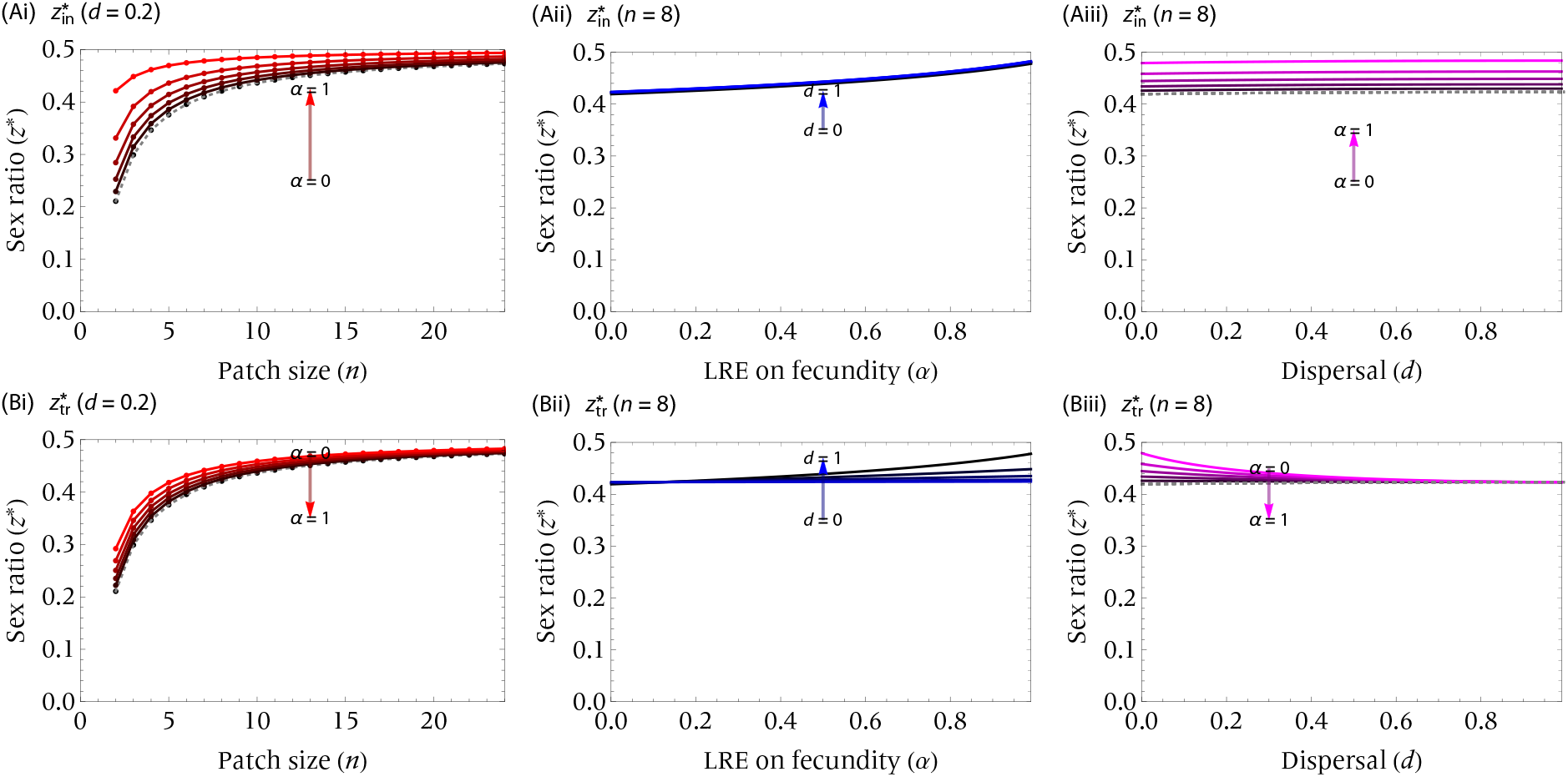
Additive male-LRE.

**SI Figure S5:**
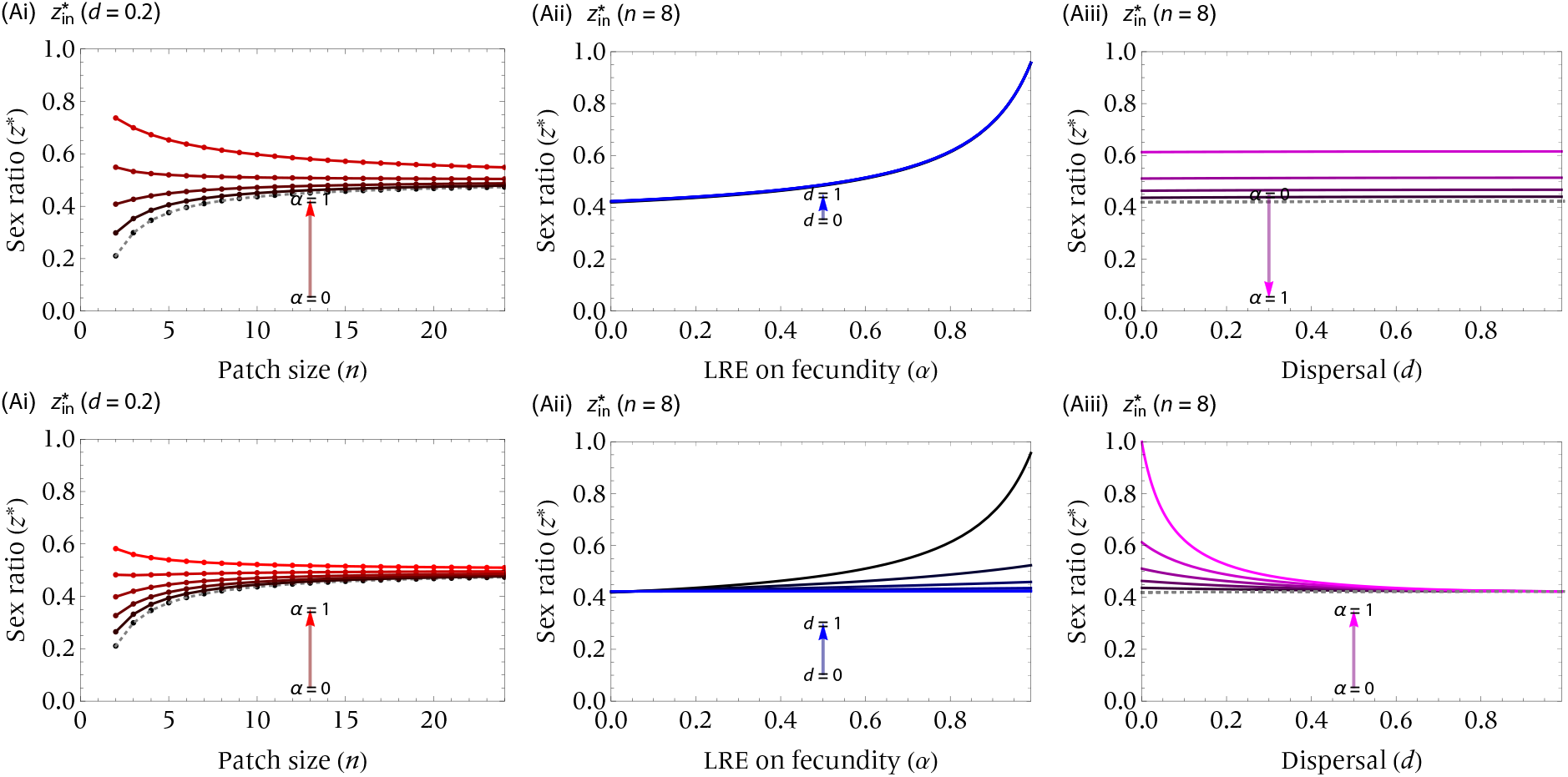
Multiplicative male-LRE. Note the difference from the previous graphic figures in the ordinate range (0 to 1).

**SI Figure 6:**
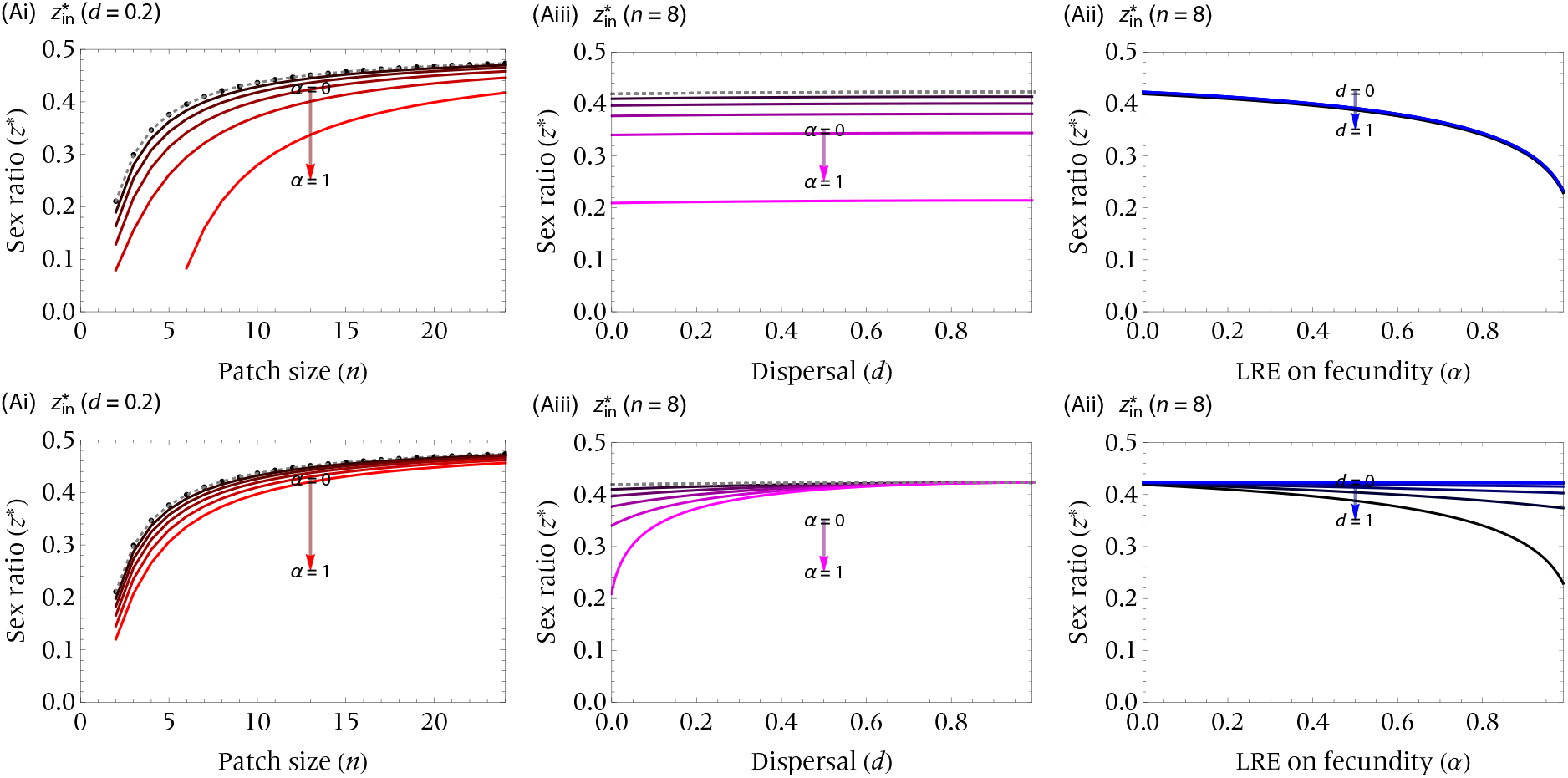
Combinational female-LRE. *θ* = *α* = 0.5. Other parameters as shown.

